# CRYPTOCHROMES confer robustness, not rhythmicity, to circadian timekeeping

**DOI:** 10.1101/2020.05.14.095968

**Authors:** Marrit Putker, David Wong, Estere Seinkmane, Nina Marie Rzechorzek, Aiwei Zeng, Nathaniel P. Hoyle, Johanna E. Chesham, Mathew D. Edwards, Kevin A. Feeney, Robin Fischer, Nicolai Peschel, Ko-Fan Chen, Christopher P. Selby, Aziz Sancar, John S. O’Neill

## Abstract

**Summary:** Circadian (approximately daily) rhythms are a pervasive property of mammalian cells, tissues, and behaviour, ensuring physiological and metabolic adaptation to solar time. Models of daily cellular timekeeping revolve around transcriptional feedback repression, whereby CLOCK and BMAL1 activate the expression of ‘clock proteins’ PERIOD (PER) and CRYPTOCHROME (CRY), which in turn repress CLOCK/BMAL1 activity. CRY proteins are thus considered essential negative regulators of the oscillation; a function supported by behavioural arrhythmicity of CRY-deficient mice when kept under constant conditions. Challenging this interpretation, however, we find evidence for persistent circadian rhythms in mouse behaviour and cellular PER2 levels when CRY is absent. CRY-less oscillations are variable in their expression and have a shorter period than wild type controls. Importantly, we find classic circadian hallmarks such as temperature compensation and determination of period by casein kinase 1δ/ε activity to be maintained. In the absence of CRY-mediated transcriptional feedback repression and rhythmic *Per2* transcription, PER2 protein rhythms are sustained for several cycles, accompanied by circadian variation in protein stability. We suggest that, whereas circadian transcriptional feedback imparts robustness and functionality onto biological clocks, the core timekeeping mechanism is post-translational. Our findings suggest that PER proteins normally act as signalling hubs that transduce timing information to the nucleus, imparting daily rhythms upon the activity of transcriptional effectors.

**Highlights:**

➢ PER/CRY-mediated negative feedback is dispensable for mammalian circadian timekeeping
➢ Circadian variation in PER2 levels persists in the absence of rhythmic *Per2* transcription
➢ CK1 and GSK3 are plausible mechanistic components of a ‘cytoscillator’ mechanism
➢ CRY-mediated feedback repression imparts robustness to biological timekeeping

**In brief:** Circadian turnover of mammalian clock protein PERIOD2 persists in the absence of canonical transcriptional feedback repression and rhythmic clock gene activity, demanding a re-evaluation of cellular clock function and evolution.

## Introduction

The adaptive advantage conferred on organisms by anticipation of the 24-hour cycle of day and night has selected for the evolution of circadian clocks that, albeit in different molecular forms, are present throughout all kingdoms of life (Edgar et al., 2012; Rosbash, 2009). Circadian rhythms are robust, in that they are “capable of performing without failure under a wide range of conditions” (Merriam-Webster Dictionary, 2020). The mechanism proposed to generate daily timekeeping in mammalian cells is a delayed transcriptional-translational feedback loop (TTFL) that consists of activating transcription factor complexes containing CLOCK and BMAL1 and repressive complexes, containing the BMAL1:CLOCK targets PERIOD and CRYPTOCHROME (Dunlap, 1999; Reppert and Weaver, 2002; Takahashi, 2016). Various coupled, but non-essential, auxiliary transcriptional feedback mechanisms are thought to fine-tune the core TTFL and coordinate cell-type specific temporal organisation of gene expression programs; the best characterised being effected by the E-box mediated rhythmic expression of REV-ERBα/β, encoded by the Nr1d1/2 genes ((Liu et al., 2008; Preitner et al., 2002; Takahashi, 2016; Ueda, 2007). These auxiliary loops are not considered sufficient to generate circadian rhythms in the absence of the core TTFL (Liu et al., 2008; Preitner et al., 2002).

CRY1 and CRY2 operate semi-redundantly as the essential repressors of CLOCK/BMAL1 activity (Chiou et al., 2016; Ye et al., 2014), required for the nuclear import of PER proteins, and together are considered indispensable for circadian regulation of gene expression *in vivo* as well as in cells and tissues cultured *ex vivo* (Chiou et al., 2016; Kume et al., 1999; Ode et al., 2017; Sato et al., 2006). Certainly, mice homozygous null for *Cry1* and *Cry2* do not express circadian behavioural rest/activity cycles under standard experimental conditions (Horst and Muijtjens, 1999; Thresher et al., 1998; Vitaterna et al., 1999).

The hypothalamic suprachiasmatic nucleus (SCN) is a central locus for circadian coordination of behaviour and physiology, and research over the last two decades has stressed the strong correlation between SCN timekeeping *in vivo* and its activity when cultured *ex vivo* (Anand et al., 2013; Welsh et al., 2010). We were therefore intrigued by the observation that roughly half of organotypic SCN slices prepared from homozygous *Cry1-/-,Cry2-/-* (CRY knockout; CKO) mouse neonates continue to exhibit approximately short period ~20h-hour rhythms, observed using the genetically encoded PER2::LUC clock protein::luciferase fusion reporter (Maywood et al., 2011; Ono et al., 2013), despite having previously been described as arrhythmic (Liu et al., 2007). Moreover, short period circadian rhythms of locomotor activity have previously been reported for CKO mice raised from birth under constant light (Ono et al., 2013). As CKO SCN oscillations were only observed in cultured neonatal organotypic slices *ex vivo,* they were suggested to be a network-level SCN-specific rescue by the activity of neuronal circuits, that desynchronise during post-natal development (Ono et al., 2013; Welsh et al., 2010). In our view, however, these observations are difficult to reconcile with an essential requirement for CRY in the generation of circadian rhythms. Rather, they are more consistent with CRY making an important contribution to circadian rhythm stability and functional outputs, as recently shown for the genes *Bmal1* and *Clock* (Landgraf et al., 2016; Ray et al., 2020), which had both previously been thought indispensable for circadian timekeeping in individual cells (Bunger et al., 2000; DeBruyne et al., 2007).

Recent observations have further questioned the need for transcriptional feedback repression to enable cellular circadian timekeeping. For example, circadian protein translation is regulated by cytosolic BMAL1 through a transcription-independent mechanism (Lipton et al., 2015), and isolated erythrocytes exhibit circadian rhythms despite lacking any DNA (Cho et al., 2014; O’Neill and Reddy, 2011). Moreover, circadian timekeeping in some species of eukaryotic alga and prokaryotic cyanobacteria can occur entirely post-translationally (Nakajima et al., 2005; O’Neill et al., 2011; Sweeney and Haxo, 1961; Tomita et al., 2005). Whether non-transcriptional clock mechanisms operate in other (nucleated) mammalian cells is unknown however, and hence their mechanism and relationship with TTFL-mediated rhythms is an open question.

Here, we used cells and tissues from CRY-deficient mice, widely accepted not to exhibit circadian transcriptional regulation (Edwards et al., 2016; Kume et al., 1999; Ukai-Tadenuma et al., 2011) to test whether any timekeeping function remained from which we might begin to dissect the mechanism of the postulated transcription-independent cytosolic oscillator, or ‘cytoscillator’ (Hastings et al., 2008).

## Results

### Cell-autonomous circadian PER2::LUC rhythms in the absence of CRY proteins

Consistent with previous observations, we observed no significant circadian organisation of locomotor activity in CRY-deficient (CKO) mice following entrainment to 12h:12h light:dark (LD) cycles or in constant light (LL). Upon transition from constant light to constant darkness (DD) (described to be a stronger zeitgeber (Chen et al., 2008)) however, CKO mice expressed rhythmic bouts of consolidated locomotor activity with a period of ~16.5h (Figure 1A, B, S1A). CKO rhythms were shorter in period and more variable than wild type (WT) controls, but persisted for >2 weeks, consistent with these mice possessing residual timing function that is not engaged during standard environmental entrainment protocols. In support of this interpretation, and in accordance with previous reports (Maywood et al., 2011; Ono et al., 2013), longitudinal bioluminescence recordings of organotypic PER2::LUC SCN slices cultured *ex vivo* from WT or CKO neonates revealed rhythmic PER2 expression in approximately 40% of CKO slices (Figure 1C). In line with behavioural data and previous reports, these CKO SCN rhythms exhibited significantly shorter periods compared with WT controls (Figure 1D).

**FIGURE 1.**
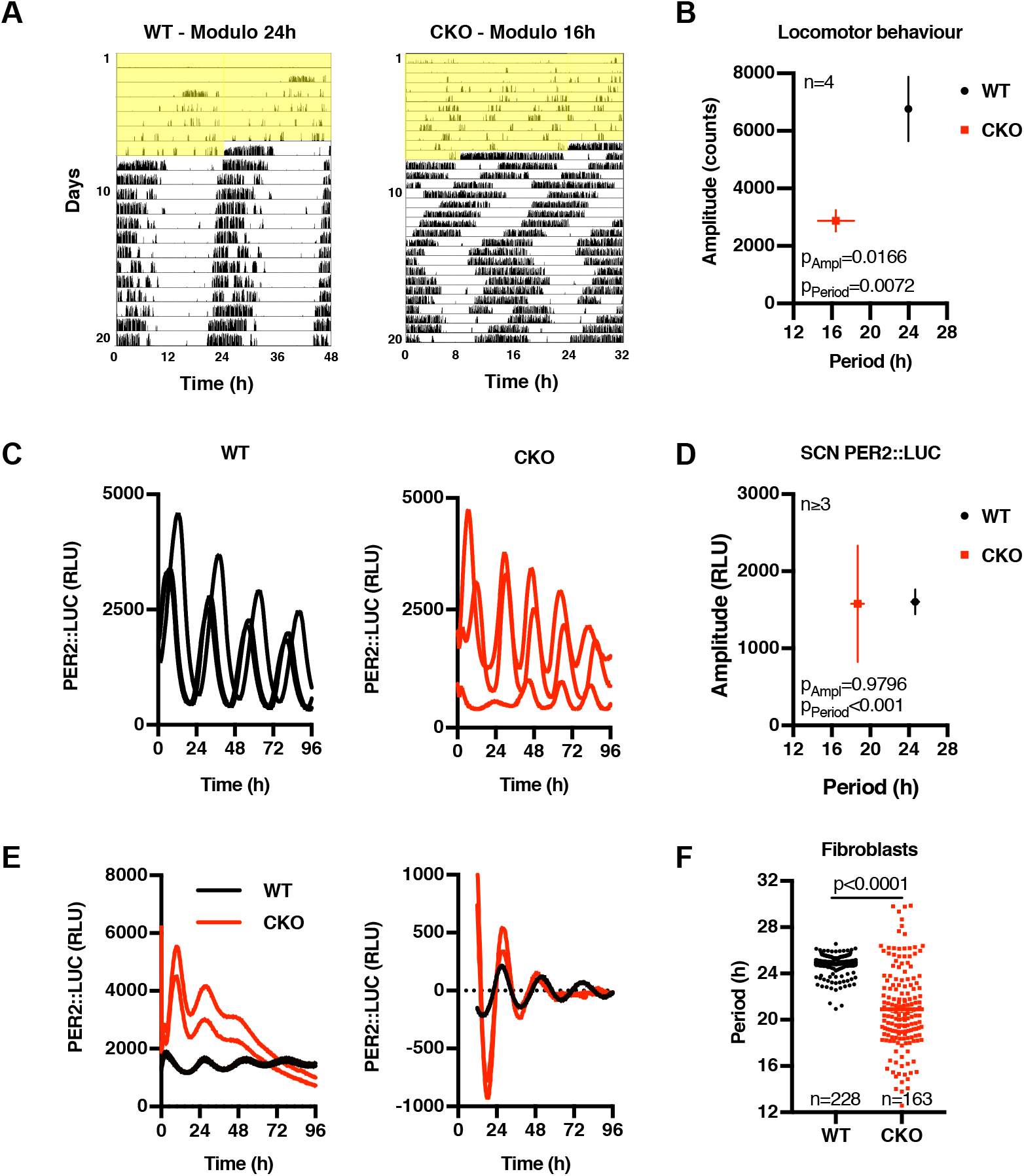
CRY-independent circadian timekeeping occurs cell-autonomously. (A) Representative double-plotted actograms showing wheel-running activity of wild type (WT) and CRY-deficient (CRY Knockout; CKO) mice during constant light (yellow shading) and thereafter in constant darkness. Note the 48 hour X-axis for WT versus 32 hour for CKO. (B) Mean period and amplitude (±SEM) of mouse behavioural data (n=4). (C) Longitudinal bioluminescence recordings of organotypic SCN slices from WT (black) and CKO (red) PER2::LUC mice (RLU; relative light units). (D) Mean period and amplitude (±SEM) of rhythmic SCN bioluminescence traces. (E) Circadian PER2::LUC expression in immortalised WT and CKO adult lung fibroblasts. Left panel shows two raw traces of a representative longitudinal bioluminescence recording, right panel shows same data detrended with a 24-hour moving average to remove differences in baseline expression. (F) Period of rhythmic fibroblast bioluminescence traces from at least 31 experiments (n≥3 per experiment). P-values were calculated using an unpaired t test with Welch correction. Standard deviations differ significantly between WT and CKO (F test: p <0.0001).

Two explanations might account for the variable CKO SCN phenotype: (1) the previously proposed explanation: genetic loss-of-function is compensated at a network-level by SCN-specific neuronal circuits whose function is sensitive to developmental phase and small variations in slice preparation (Evans et al., 2012; Liu et al., 2007; Ono et al., 2013; Tokuda et al., 2015); or (2) CKO (SCN) cells have cell-intrinsic circadian rhythms that are expressed (or observed) more stochastically and with less robustness than their WT counterparts, and can be amplified by SCN interneuronal signalling (O’Neill and Reddy, 2012; Welsh et al., 2010).

To distinguish between these two possibilities, we asked whether PER2::LUC rhythms are observed in populations of immortalised PER2::LUC CKO adult fibroblasts, which lack the specialised interneuronal neuropetidergic signalling that is so essential to SCN amplitude and robustness *in* and *ex vivo* (O’Neill and Reddy, 2012; Welsh et al., 2010). We observed this to be the case (Figure 1E and S1B-C). Across >100 recordings, using independently-generated cell lines cultured from multiple CRY-deficient mice (male and female), we observed PER2::LUC rhythms that persisted for several days under constant conditions. Again, the mean period of rhythms in CRY-deficient cells was significantly shorter than WT controls, and with increased variance within and between experiments (F-test p-value <0.0001, Figure 1F and S1D, E). Consistent with SCN results, rhythmic PER2::LUC expression in CKO cells occurred stochastically between experiments, being observed in ~30% of independently performed assays. Importantly, there was very little variation in the occurrence of rhythmicity within experiments meaning that in any given recording all CKO replicate cultures were rhythmic or none, whereas WT cultures were always rhythmic. CKO PER2::LUC rhythms damped more rapidly than wild type controls (Figure S1F), and were more sensitive to acute changes in temperature than WT controls (Figure 2A, C), consistent with their oscillation being less robust. Crucially though, the PER2::LUC rhythms in CKO cells were temperature compensated (Figure 2A, B) and entrained to 12h:12h 32°C:37°C temperature cycles in the same phase as WT controls (Figures 2C), and are thus circadian by definition (Pittendrigh, 1960).

**FIGURE 2.**
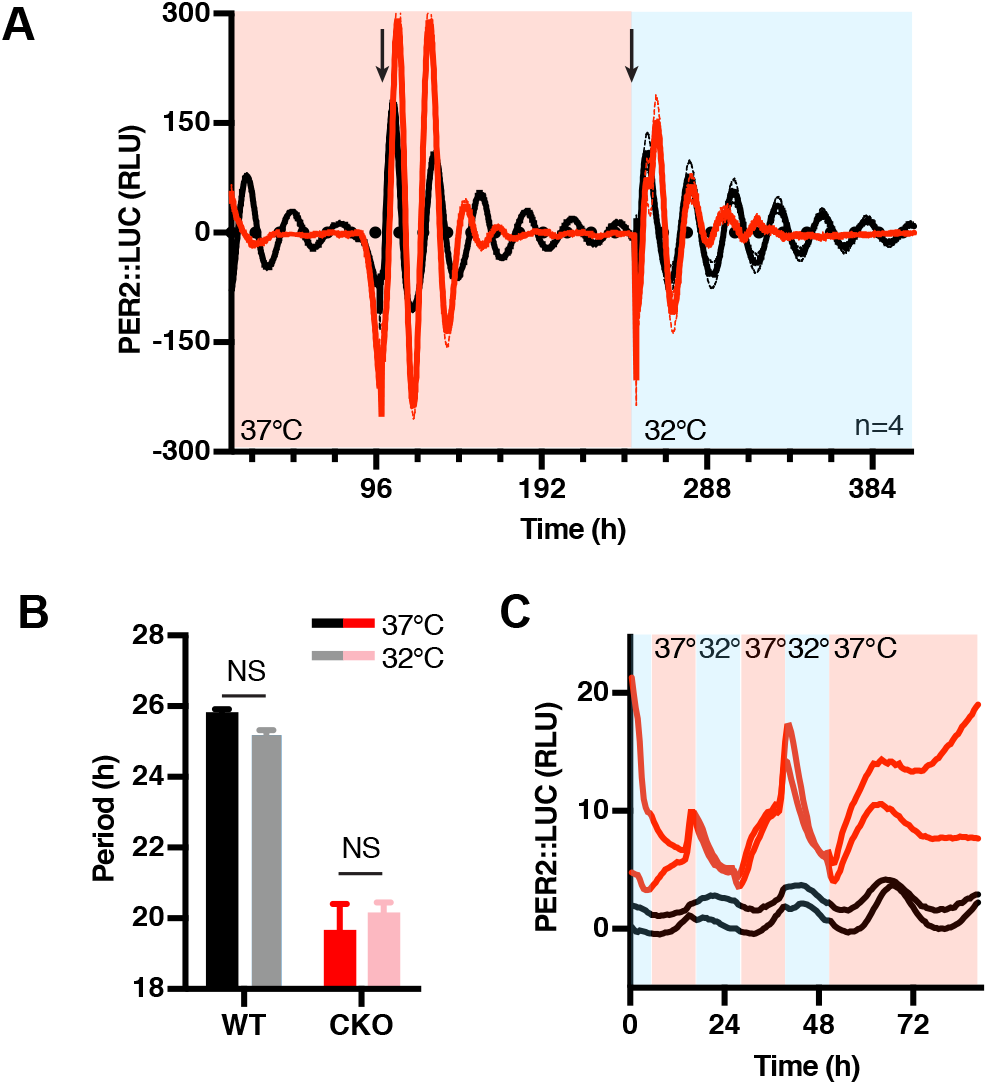
CRY-less oscillations are temperature compensated and entrained. (A) Detrended traces of bioluminescence recordings of WT and CKO fibroblast at different constant temperature conditions within the physiological range (n=4, mean ± SEM). Temperature was changed from 37°C to 32°C halfway through the experiment, as depicted by red/blue shading. Arrows represent medium changes. Note the lack of rhythmicity in the first three days in CKO and the appearance of rhythmicity after the first medium change. (B) Quantification of period from recordings presented in (A). Both WT and CKO oscillations are temperature compensated with respective Q_10_s of 1.05 and 0.95. (C) Bioluminescence of WT and CKO PER2::LUC cells during temperature entrainment (12h 32°C (blue) – 12h 37°C (red)). Two representative traces of two independent cell lines are shown per genotype.

These observations suggest that CRY-dependent transcriptional feedback repression primarily confers robustness to rhythmic clock output, rather than generating circadian rhythms *per se.* To test this in another model system we turned to *Drosophila melanogaster,* where TIMELESS fulfils the functionally analogous role to mammalian CRY proteins as the obligate partner of PER, required for repression of circadian transcription at E-box promotor elements. In assays of PER:LUC (XLG-LUC) bioluminescence in freely behaving flies, we observed robust circadian rhythms in 7 out of 36 TIMELESS knockout animals, compared with 11 out of 21 wild type controls (Figure S2). As observed for CRY-deficient cells, rhythms in TIMELESS-deficient flies persisted over several days, but were noisier and exhibited lower relative amplitude than WT.

Considering recent reports that transcriptional feedback repression is not absolutely required for circadian rhythms in the activity of FRQ, the functional orthologue of PER in the fungus *Neurospora crassa* (Larrondo et al., 2015), that nascent transcription is not required for circadian rhythms in the green lineage (O’Neill et al., 2011), or in isolated human red blood cells (O’Neill and Reddy, 2011), we next asked whether residual rhythms of PER2:LUC in CRY-deficient cells result from post-translational regulation.

### CRY-independent PER2::LUC rhythms are driven by a non-transcriptional process

CRY has previously been described as the driving factor for feedback repression of BMAL1/CLOCK-dependent transcriptional activation, and is therefore considered essential to the rhythmic regulation of clock-controlled genes (CCGs). In fact, overexpression studies have suggested PER requires CRY to exert its function as a BMAL1-CLOCK repressor (Chiou et al., 2016; Ye et al., 2014). This importance of CRY for BMAL1-CLOCK repression (and auto-repression of *Cry* and *Per*) was also suggested by the increased PER2::LUC levels observed in CKO cells (Figure 1E, S1B). Indeed, at the peak of PER2::LUC expression, CKO cells contain approximately twice as many PER2 molecules compared with their WT counterparts (Figure 3A and S3A).

**FIGURE 3.**
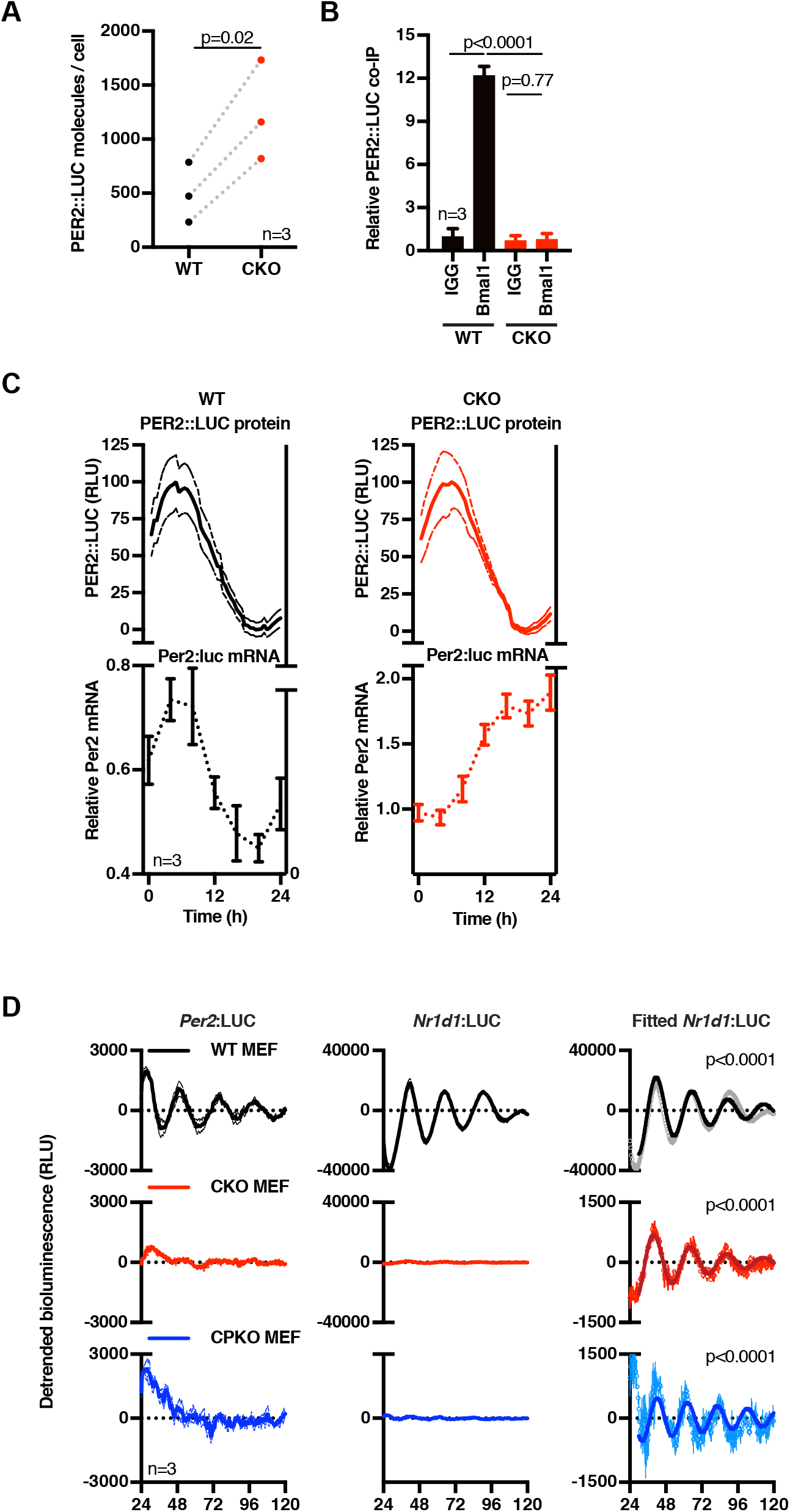
CRY-independent rhythms are regulated post-transcriptionally. (A) Mean number of PER2::LUC molecules per cell at the estimated peak of PER2 expression for each cell line (mean of three experiments, n=3 each). P-values were calculated in a paired t test. (B) PER2::LUC binding to BMAL1 in WT and CKO cells. Cells were harvested at the peak of PER2 expression, BMAL1 was immunoprecipitated, and PER2::LUC binding was measured by bioluminescence measurements (n=3, mean ±SD). P-values were calculated in an unpaired t test. (C) Per2 mRNA levels in WT (left) and CKO (right) cells were determined by qPCR over one circadian cycle (bottom), while PER2::LUC bioluminescence (min-max normalised) was recorded from parallel cultures (top) (mean Per2 mRNA relative to Rns18s (bottom) and PER2::LUC signal (top), n=3, ±SEM). The WT mRNA trace could be fitted with a circadian damped sine wave (p=0.0412) whereas data of CKO cells could not (ns). (D) Detrended *Per2* and *Nr1d1* promoter activity in WT, CKO and quadruple *Cry1/2-Per1/2* knockout (CPKO) mouse embryonic fibroblasts (MEFs) recorded at 37°C. *Nr1d1* data were fit with a circadian damped sine wave over straight line (p<0.0001) (right graphs). Similar recordings performed at 32°C and an expanded view of *Per2* data are in figures S3E and F.

Although not sufficient to completely rescue rhythms in CKO cells, it seemed plausible that increased PER protein expression might partially compensate for the loss of CRY function and continue to exert auto-regulation through rhythmic BMAL1-CLOCK binding, thereby accounting for the residual PER2::LUC rhythms in CKO cells. To test this possibility, we compared BMAL1-PER2 binding at the expected peak of BMAL1-PER2 complex formation (i.e. at the peak of PER2::LUC expression) in WT and CKO cells. To this end, we immunoprecipitated BMAL1 and measured the associated PER2::LUC activity. In accordance with CRY being required for PER2-BMAL1 binding, we did not find a PER2::LUC-BMAL1 complex in CKO cells, while the complex was readily detected in WT cells (Figure 3B and S3B), strongly suggesting that residual oscillations in PER2::LUC cannot result from a residual negative feedback upon the BMAL1-CLOCK complex.

In the absence of PER:CRY-mediated feedback repression, it seemed unlikely that CRY-independent oscillations in PER2::LUC expression are driven directly by rhythms in *Per2* transcription. Indeed, whereas PER2::LUC in co-recorded cells showed a clear variation over 24h, *Per2* mRNA in parallel replicate CKO cultures instead exhibited a gradual accumulation (Figure 3C). In contrast and as expected, *Per2* mRNA in WT cells varied in phase with co-recorded PER2::LUC oscillations. The gradual increase of *Per2* mRNA in CKO cells is concordant with *Per2* transcriptional derepression predicted by the canonical TTFL model, accounting for the generally increased levels of PER2::LUC we observed (Figure 3A), but not their oscillation. In agreement with these findings and in contrast with WT cells, *Bmal1* mRNA also showed no significant variation in CKO cells (Figure S3C), suggesting that E-box-dependent circadian regulation of REV-ERB activity may not occur in the absence of CRY-mediated feedback repression. In an independent validation we assessed the activity of the circadian E-box-driven Cry1-promoter (Maywood et al., 2013) in mouse adult WT and CKO lung fibroblasts (MAFs) (Figure S3D), as well as the *Per2-* and *Rev-erb**α***-(*Nr1d1-*) promoters in mouse embryonic fibroblasts (MEFs) (Figure 3D, S3E-G). No rhythmic *Cry1-* or *Per2-* promoter activity was observed in either set of CKO cells under any condition, whereas isogenic control cells showed clear circadian regulation of these promoters.

In recordings from *Nr1d1:LUC* MEFs however, we were most surprised to observe temperature-compensated circadian rhythms in the activity of the Nr1d1 promoter in CKO cells, at just ~3% amplitude of WT cells, that persisted for several days (Figure 3D and S3E, red traces). In the same experiments, similar but still noisier and lower amplitude rhythms were also detected in quadruple knockout MEFs that were also deficient for PER1/2, as well as CRY1/2 (CPKO, Figure 3D and S3E, blue traces), confirming these oscillations cannot be attributable to any vestigial activity of PER proteins. We acknowledge it is conceivable that some unknown TTFL-type mechanism might generate these residual oscillations in *Nr1d1* promoter activity. However, we find it more plausible that residual oscillations of *Nr1d1*:LUC in CKO cells are the output of a post-translational timekeeping mechanism, from which the amplification and robustness conferred by CRY-dependent transcriptional feedback repression has been subtracted. Indeed, we note that besides CRY, *Nr1d1* expression is regulated by many other transcription factors, e.g., AP-1, NRF2, NF-KB and BMAL1/CLOCK (Preitner et al., 2002; Wible et al., 2018; Yang et al., 2014), whose activity is regulated post-translationally by the same rather promiscuous kinases that rhythmically regulate PER and BMAL1 in other contexts (Eide et al., 2002; Iitaka et al., 2005; Narasimamurthy et al., 2018; Sahar et al., 2010) e.g. casein kinase 1, glycogen synthase kinase (Jiang et al., 2018; Liang and Chuang, 2006; Medunjanin et al., 2016; Preitner et al., 2002; Rada et al., 2011; Tullai et al., 2011).

### Circadian control of PER2 stability persists in absence of CRY

The concentrations of luciferase substrates (Mg.ATP, luciferin, O_2_) under our assay conditions are >10x higher than their respective K_m_ (Feeney et al., 2016a) and so it is implausible that PER2::LUC rhythms in CKO cells result from anything other than circadian regulation in the abundance of the PER2::LUC fusion protein. Indeed, PER2::LUC levels measured in cell lysates perfectly mirrored longitudinal PER2::LUC recordings from both WT and CKO cells (Figure 4A). We observed that the addition of the proteasomal inhibitor MG132 to asynchronous cells led to acute increases in PER2::LUC levels which were significantly greater in CKO cells than in WT controls, indicating that CKO cells support higher basal rates of PER2 turnover (Figure 4B and 4C). In consequence therefore, relatively small changes in the rate of PER2::LUC translation or degradation should be sufficient to affect the steady state PER2::LUC concentration. CKO cells exhibit no rhythm in *Per2* mRNA (Figure 3C, D), nor do they show a rhythm in global translational rate (Figure S4A, B), nor did we observe any interaction between BMAL1 and S6K/eIF4 as occurs in WT cells (Lipton et al, 2015) (Figure S4C). We therefore investigated whether changes in PER2::LUC stability might be responsible for the persistent bioluminescence rhythms in CKO cells, by analysing the decay kinetics of luciferase activity during saturating translational inhibition.

**FIGURE 4.**
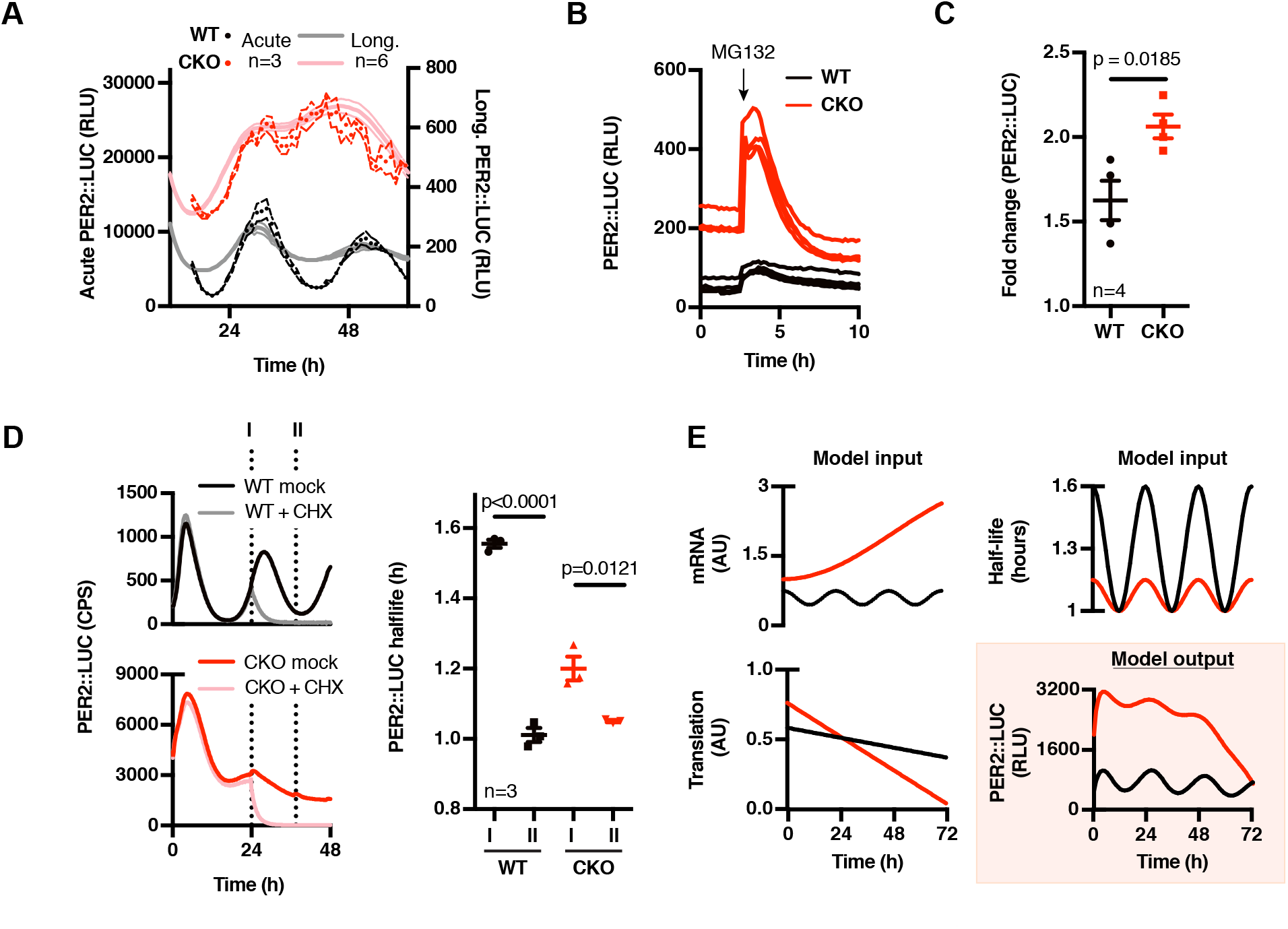
PER2::LUC stability oscillates in CRY-deficient cells. (A) Actual PER2::LUC levels (dark symbols (3-hour moving average, n=3 ±SEM, 4 outliers removed)) as assayed in acute luciferase assays on cell lysates from cells harvested every hour over 48 hours, compared with parallel longitudinal co-recordings from cells in the presence of 0.1mM luciferin (light lines (n=6, mean ±SEM)). (B) PER2::LUC recording of asynchronous WT and CKO cells pulsed with proteasome inhibitor MG132 (10 μM, applied at the arrow) (n=3, mean ±SEM). (C) Quantification of relative PER2::LUC induction upon proteasome inhibition. P-value was calculated by unpaired t test. (D) Phase-dependent PER2::LUC half-life was determined by inhibiting translation at different circadian phases and fitting the resulting data with a one-phase exponential decay curve (n=3, mean ±SEM). Left image depicts the timing of cycloheximide (CHX, 10 μM) pulses (labelled I (PER2 levels going up) and II (PER2 levels going down)), plotted on PER2::LUC bioluminescence traces of control cells (dark colours). A representative trace of CHX-treated cells at time point I is shown in light colours. See FIG S4D-E for more raw data and time points. Right image shows quantifications, p-values were calculated by unpaired t test. (E) A simple model incorporating mRNA, protein translation and PER2::LUC stability we measured experimentally (inputs) shows that the observed oscillating stability of PER2 is sufficient to generate rhythmic PER2::LUC expression (output).

In the presence of 10 μM cycloheximide (CHX) PER2::LUC bioluminescence decayed exponentially (Figure 4D and S4D, R2>0.9), with a half-life that was consistently <2 hours (Figure 4D and S4E); much less than the half-life of luciferase expressed in fibroblasts under a constitutive promoter (≥5 h, Figure S4D and E). Moreover, we observed a significant variation (±50%) in the half-life of PER2::LUC between the rising and falling phases of its expression (1.5 vs 1 h, respectively, Figure 4D and S4F) without any commensurate change in global protein turnover (Figure S4G). Strikingly, we also observed a similar phase-dependent variation of PER2::LUC stability in CKO cells, with a smaller (±20%) but significant difference between opposite phases of the oscillation (Figure 4D). To test if a 20% variation in protein half-life, in the absence of any underlying mRNA abundance rhythm, was sufficient to account for our experimental observations given the intrinsically high turnover of PER2, we made a simple mathematical model using experimentally derived values for mRNA level, protein half-life and translation (Figure 3C and S4). We found that the model produced PER2::LUC levels that closely approximate our experimental observations (Figure 4E). Thus whilst we cannot absolutely discount the possibility that rhythmic translation contributes to the PER2::LUC rhythms in CKO cells, we found no evidence to support this, whereas experimental observations and theoretical modelling do suggest rhythmic PER2 degradation alone is sufficient to explain the residual bioluminescence rhythms we observe in CKO PER2::LUC fibroblasts.

### CK1δ/ε and GSK3 contribute to CRY-independent PER2 oscillations

PER2 stability is primarily regulated through phosphorylation by casein kinases (CK) 1δ and 1ε, which phosphorylate PER2 at phosphodegron sites to target it for proteasomal degradation (Lee et al., 2009; Philpott et al., 2020). In this context CK1δ/ε frequently operate in tandem with glycogen synthase kinase (GSK) 3α/β, as occurs in the regulation of β-catenin stability (O’Neill et al., 2013; Robertson et al., 2018). Interestingly, both CK1δ/ε and GSK3α/β have a conserved role in determining the speed at which the eukaryotic cellular circadian clock runs (Causton et al., 2015; Hastings et al., 2008), both in the presence and absence of transcription (Beale et al., 2019; Hirota et al., 2008; Meng et al., 2008; O’Neill et al., 2011). This is despite the fact that the clock proteins phosphorylated by these kinases are highly dissimilar between animals, plants, and fungi (Causton et al., 2015; Wong and O’Neill, 2018).

We hypothesised that the PER2::LUC rhythm in CKO cells reflects the continued activity of a post-translational timekeeping mechanism that involves CK1δ/ε and GSK3α/β, which results in the differential phosphorylation and turnover of clock protein substrate effectors such as PER2 during each circadian cycle (O’Neill et al., 2013). To test this we incubated WT and CKO cells with selective pharmacological inhibitors of CK1δ/ε (PF670462; PF) and GSK3α/β (CHIR99021; CHIR), which have previously been shown to slow down, and accelerate, respectively, the speed at which the cellular clock runs in a wide range of model organisms (Badura et al., 2007; Causton et al., 2015; Hirota et al., 2008; O’Neill et al., 2011). As a control we used KL001, a small molecule inhibitor of CRY degradation (Hirota et al., 2012), which has previously been shown to affect cellular rhythms in WT cells via increased CRY stability.

We found that inhibition of CK1δ/ε and GSK3-α/β had the same effect on circadian period in CKO cells as WT controls (Figure 5A, B, S5A, B). In contrast, KL001 increased period length and reduced amplitude of PER2::LUC expression in WT cells but had no significant effect on post-translationally regulated PER2::LUC rhythms in CKO cells (Figure 5C and S5C). Besides confirming the specific mode of action for KL001 in targeting CRY stability, these observations implicate CK1δ/ε and GSK3α/β in regulating the post-translational rhythm reported by PER2::LUC in CKO cells.

**FIGURE 5.**
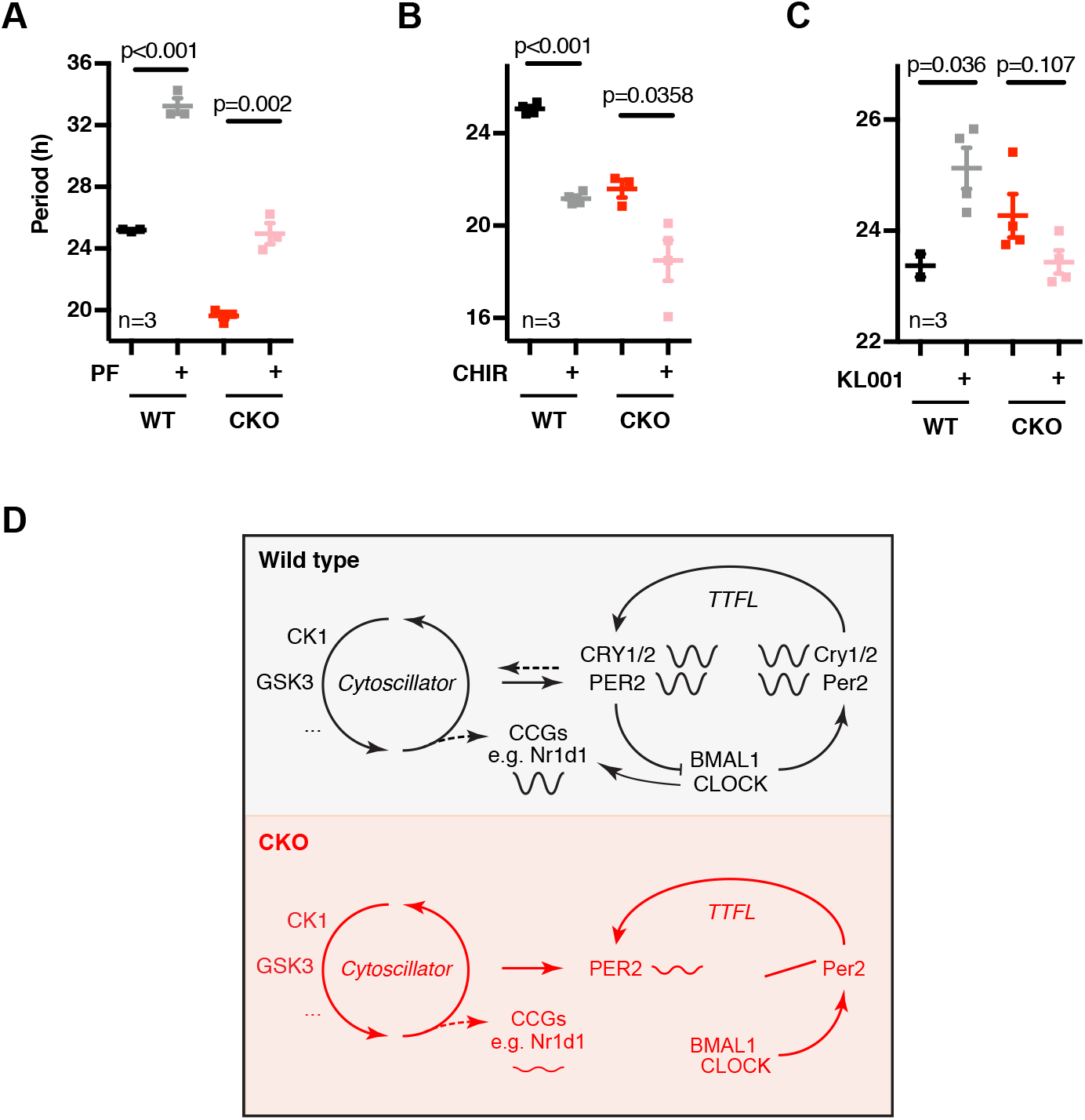
A role for CK1 and GSK3 in the cytoplasmic oscillator. (A) Period (right; n=3, mean ±SEM) analyses of WT and CKO PER2::LUC cells in the presence or absence of CK1δ/ε inhibitor PF670462 (0.3 μM; PF). P-values were calculated by unpaired t test. (B) As in (A), GSK3 inhibitor CHIR99021 (5 μM; CHIR). (C) As in (A), in presence of CRY inhibitor KL001 (1 μM). (D) Schematic model integrating CRY-independent timekeeping into the existing canonical model of the circadian clock. The CRY-dependent gene expression feedback loop (TTFL) is required for most circadian regulation of transcriptional clock controlled genes (CCGs) and therefore for robustness and behavioural and physiological rhythmicity. However, it is dispensable for circadian timekeeping *per se,* as reported by residual oscillations in PER2 protein levels, suggestive of the existence of a coupled underlying (cytosolic) timekeeping mechanism involving CK1 and GSK3 (cytoscillator). See FIG S5 for raw data.

## Discussion

We found that transcriptional feedback in the canonical TTFL clock model is dispensable for cell-autonomous circadian timekeeping in animal and cellular models. In mice and flies, deficient for CRY/PER or TIMELESS/PER-mediated feedback repression, the capacity for circadian gene expression remained intact, though clearly impaired with respect to WT. Circadian rhythms of PER2 abundance were observed in CKO SCN slices and fibroblasts, indicating that the post-translational mechanisms that confer circadian rhythmicity onto PER proteins in WT cells remain ostensibly intact in the absence of CRY-mediated transcriptional feedback repression. Importantly however, CKO PER2 rhythms were only observed in a minority of recordings (~30%), and when observed they showed increased variance of period and sensitivity to perturbation. This reduced capacity to perform without failure under a wide range of conditions means that CRY-deficient PER2 oscillations are less robust than those in WT cells (Merriam-Webster Dictionary, 2020). We were unable to identify all of the variables that contribute to the apparent stochasticity of CKO PER2::LUC oscillations, and so cannot distinguish whether this variability arises from reduced fidelity of PER2::LUC as a circadian reporter or impaired timing function in CKO cells. In consequence, we restricted our study to those recordings in which clear bioluminescence rhythms were observed, enabling the interrogation of TTFL-independent cellular timekeeping.

In the field of chronobiology, CKO cells and mice are often used as clock-deficient models. Indeed, canonical circadian transcriptional output is essentially absent from these models (Hoyle et al., 2017; Ode et al., 2017), and thus for studying TTFL-mediated control of overt physiology they are appropriate negative controls. However, as the underlying timekeeping mechanism seems at least partially intact, we therefore consider it inappropriate to describe CKO cellular models as arrhythmic. Indeed, rest/activity behaviour of CKO mice does entrain to daily cycles of restricted feeding (Iijima et al., 2005), which is SCN-independent (Storch and Weitz, 2009), as well as a sufficiently strong synchronising zeitgeber (Figure 1A, (Chen et al., 2008)). Thus, non-TTFL mediated timekeeping seems sufficient to serve as an (about) daily interval timer *in vivo* (Crosby et al., 2019).

Previous studies have reported isolated CKO cells to be entirely arrhythmic (Ode et al., 2017; Sato et al., 2006; Ukai-Tadenuma et al., 2011), in stark contradiction with our findings. However, most such studies measured changes in transcription either by quantitative RT-PCR, or with luciferase fusions to fragments of the *Bmal1, Per* and *Cry* promoters which we also found to be arrhythmic in CKO cells. We did observe low amplitude oscillations in *Nr1d1* promoter activity however. It may be pertinent to report that these MEF recordings only revealed oscillations of *Nr1d1*-promoter activity, and only in bicarbonate-buffered medium supplemented with 1mM luciferin and 10% serum (Figure 3D), but not in low serum or HEPES-buffered media, as employed in other studies that used different circadian reporters and may have employed sub-saturating concentrations of luciferin (Feeney et al., 2016a). It is also plausible that the high sensitivity of the electron-multiplying CCD camera we used for these bioluminescence assays allows the quantification of biological rhythms that were not detectable using other approaches (Crosby et al., 2017).

Although several mechanisms for circadian regulation of translation have been described (Jouffe et al., 2013; Lipton et al., 2015), we did not find any contribution of rhythmic translation to CRY-independent rhythms. In fact, the BMAL1-S6K1 interaction that mediates BMAL1’s interaction with the translational apparatus is absent from CKO cells (Figure S4C), implying a possible role for CRY proteins in this complex. Instead, we found an overt circadian regulation of PER2::LUC stability that persists in the absence of CRY proteins and which was sufficient to account for the observed PER2::LUC rhythms in a simple mathematical model. Persistent post-translational regulation of PER stability/activity may also account for the results of earlier over-expression studies, in mammalian cells and flies, where constitutive *Per* mRNA expression resulted in rhythmic PER protein abundance (Fujimoto et al., 2006; Yamamoto et al., 2005; Yang and Sehgal, 2001); whereas *Per* over-expression should really abolish rhythms if *Per* mRNA levels are the fundamental state variable of the oscillation. This interpretation has marked similarities with recent reports in the fungal clock model, *Neurospora Crassa,* where experiments have suggested that post-translationally regulated cycles in the activity of the FRQ clock protein, not its abundance, are the critical determinant of downstream circadian gene regulation (Larrondo et al., 2015).

Indeed, our observations may not be particularly surprising when one considers that post-translational regulation of circadian timekeeping is ubiquitous in eukaryotes, with the period-determining function of CK1δ/ε and GSK3α/β being conserved between the animal, plant and fungal clocks (Causton et al., 2015; Hirota et al., 2010; Lee et al., 2009; O’Neill et al., 2011; Wong and O’Neill, 2018; Yao and Shafer, 2014), despite their clock protein targets being highly dissimilar between phylogenetic kingdoms. Importantly we observed that pharmacological inhibition of these kinases elicited the same period-lengthening and -shortening effects on CRY-independent rhythms as on WT rhythms. This has implications for our understanding of the role that these kinases play in the cellular clock mechanism, since in the absence of TTFL-mediated timekeeping their effects cannot be executed through regulation of any known transcriptional clock component.

Given similar findings across a range of model systems, including isolated red blood cells (Wong and O’Neill, 2018), the simplest interpretation of our findings entails an underlying, evolutionarily-conserved post-translational timekeeping mechanism: a “cytoscillator” (Hastings et al., 2008)) that involves CK1δ/ε and GSK3α/β, and can function independently of canonical clock proteins, but normally reciprocally regulates with cycles of clock protein activity through changes in gene expression (Qin et al., 2015). This cytoscillator confers 24-hour periodicity upon the activity and stability of PER2, and most likely to other clock protein transcription factors as well (Figure 3D). However, a purely post-translational timing mechanism should be rather sensitive to environmental perturbations and biological noise (Ladbury and Arold, 2012), as seen for CKO cells. Due to the geometric nature of their underlying oscillatory mechanism, relaxation oscillators are known to be particularly insensitive to external perturbations and prevalent in noisy biological systems (Muratov and Vanden-Eijnden, 2008). We therefore suggest that in wild type cells, low amplitude, cytoscillator-driven circadian cycles of clock protein activity are coupled with, reinforced and amplified by a damped TTFL-based relaxation oscillation of stochastic frequency (Chickarmane et al., 2007), resulting in high-amplitude, sustained circadian rhythms in both clock and clock-controlled gene expression. Indeed, mathematical modelling shows that such coupling can both drive the emergence of sustained oscillations in overdamped systems (In et al., 2003) and play an important role in maintaining robust oscillations in a random environment (Medvedev, 2010). This model is consistent with recent observations in the clocks of the prokaryotic cyanobacterium *Synechoccocus elongatus* (Qin et al., 2010; Teng et al., 2013) as well as the fungus *Neurospora crassa* (Larrondo et al., 2015), and the alga *Ostreococcus tauri* (Feeney et al., 2016b) (see supplementary information for an extended discussion).

Interestingly, the concept of the eukaryotic post-translational clock mechanism we propose is not new (Jolley et al., 2012; Merrow et al., 2006; Qin et al., 2010; Roenneberg and Merrow, 1998) and resembles the KaiA/B/C mechanism elucidated in cyanobacteria (Nakajima et al., 2005; Teng et al., 2013). The challenge will now be to identify additional factors that, in concert with CK1 and GSK3, and protein phosphatase 1 (Lee et al., 2011), serve as the functional equivalents of KaiA/B/C; allowing reconstitution of the mammalian circadian clock *in vitro*. (Millius et al., 2019; Nakajima et al., 2005).

Here we have uncovered PER2 as a node of interaction between a putative cytoscillator mechanism and the canonical circadian TTFL (Figure 5D). It is unlikely however that PER2 is the only interaction between the two, as *Per2-/-* knockout cells and mice exhibit competent circadian timekeeping (Xu et al., 2007), suggesting redundancy in this respect. Indeed, the residual noisy but rhythmic activity of the *Nr1d1*-promoter in the absence of both PER1/2 and CRY1/2 (Figure 3D), suggests another point-of-connection between the cytoscillator and TTFL. Moreover, both CK1 and GSK have been implicated in the phosphorylation and regulation of many other clock proteins (See table S2 in (Causton et al., 2015), also reviewed in (O’Neill et al., 2013)). Some or all of these targets may play a role in coupling the cytoscillator with TTFL-mediated clock output. We believe that it is now imperative to delineate the specific means by which the TTFL couples with the cytoscillator to effect changes in circadian phase in order that the two resonate with a common frequency.

## Conclusion

Whilst the contribution of clock protein transcription factors to the temporal coordination of gene expression, physiology and behaviour is unambiguous, the primacy of transcriptional feedback repression as the ultimate arbiter of circadian periodicity within eukaryotic cells is not. Similar to the conserved kinase-dependent regulation of the cell division cycle, we suggest the circadian cycle in diverse eukaryotes is conserved from a common ancestor, with diverse TTFL components having been recruited throughout speciation to impart robustness, signal amplification and functional specificity to the oscillation.

### Experimental procedures

Reagents were obtained from sigma unless stated otherwise. More detailed experimental procedures can be found in the supplementary information (SI).

### Mouse work

All animal work was licensed under the UK Animals (Scientific Procedures) Act 1986, with Local Ethical Review by the Medical Research Council. *Cry1/2-null* mice were kindly provided by G. T. van der Horst (Erasmus MC, Rotterdam, The Netherlands) (Horst and Muijtjens, 1999), PER2::LUC mice by J. S. Takahashi (UT Southwestern, USA) (Yoo et al., 2004) and *Cry1:LUC* mice by M. Hastings (MRC LMB, Cambridge, UK) (Maywood et al., 2013). All lines were maintained on a C57BL/6J background. For mouse behavioral studies, CKO PER2::LUC female mice aged 2-5 months, and age-matched PER2::LUC controls, were singly housed in running wheel cages with circadian cabinets (Actimetrics). They were then subject to 7 days 12h:12h LD cycles or 7 days constant light (400 lux), and then maintained in constant darkness with weekly water and food changes. Locomotor activity was recorded using running wheel activity and passive infrared detection, which was analysed using the periodogram function of ClockLab (Actimetrics). SCN organotypic slices from 7-10 day old pups were prepared as previously described (Hastings et al., 2005), and bioluminescence recorded using photomultiplier tubes (Hamamatsu).

### Mammalian cell culture

Primary fibroblasts were isolated from lung tissue (Seluanov et al., 2010) of adult wild type (WT) and *Cry1*-/-, *Cry2*-/- (CKO) PER2::LUC male and female mice, and WT and CKO *Cry1*:LUC mice. Stable WT, CKO and *Cry1*-/-,*Cry2*-/-, *Per1-/-, Per2-/-* (CPKO) mouse embryonic fibroblasts (MEFs) expressing transcriptional luciferase reporters for clock gene activity were generated by puromycin selection and cultured as described previously (Valekunja et al., 2013). MEFs were seeded into 96-well white plates at 104 cells/well and grown to confluency for 5 days under temperature cycles (12h:12h, 32°:37°) to synchronise circadian rhythms. Primary fibroblasts were cultured as described previously (O’Neill and Hastings, 2008) and immortalised by serial passage (Xu, 2005). CRY deficiency was confirmed by PCR (see SI) and Western blotting (guinea pig-anti-CRY1 and CRY2 antibodies (Lamia et al., 2011)). NIH3T3 fibroblasts expressing SV40::LUC have been described before (Feeney et al., 2016a).

### Luciferase recordings

Fibroblast recordings were performed in air medium (either HEPES or MOPS buffered (20mM), either in airtight sealed dishes (in non-humidified conditions) or open in humidified conditions (0% CO2). Air medium stock was prepared as described previously (O’Neill and Hastings, 2008) and supplemented with 2% B-27 (Life Technologies, 50X), 1 mM luciferin (Biosynth AG), 1X glutamax (Life Technologies), 100 units/ml penicillin/100 μg/ml streptomycin, and 1% FetalCloneTM III serum (HyCloneTM). Final osmolarity was adjusted to 350 mOsm with NaCl. Recordings were preceded by appropriate synchronisation (see SI for details) in presence of 0.3 mM luciferin to prevent artificially high bioluminescence activity at the start of the recording, and started immediately after a medium change from culture medium into air medium. The presented MEF recordings were performed in an ALLIGATOR (Crosby et al., 2017), and employed bicarbonate-buffered Dulbecco’s Modified Eagle Medium (10569010) with penicillin/streptomycin and 1 mM luciferin in a humidified incubator at 5% CO2, also supplemented with 2% B-27 and 10% FetalCloneTM III serum. A range of other media conditions were explored but did not produce detectable bioluminescence rhythms in CKO or CPKO cells (not shown). For pharmacological perturbation experiments (unless stated otherwise in the text) cells were changed into drug-containing air medium from the start of the recording. Mock-treatments were carried out with DMSO or ethanol as appropriate.

Bioluminescence recordings were performed in a lumicycle (Actimetrics), a LB962 plate reader (Berthold technologies) or an ALLIGATOR (Cairn Research). Acute luciferase assays were performed using a Spark 10M microplate reader (Tecan).

### Biochemistry

The number of PER2 molecules was determined by harvesting a known number of synchronised WT and CKO cells at the peak of PER2 expression and comparing the Luciferase activity to a standard curve of recombinant Luciferase (see SI for details). Three technical replicates were measured in every experiment and the experiment was carried out three times. A representative experiment is shown.

For determining *Per2::Luc* and *Bmal1* mRNA levels, synchronised cells were harvested from constant conditions in triplicate every four hours from 24 hours up to 48 hours after media change. RNA extraction and qPCR were performed as detailed in the SI. Analysis involved three technical and three biological replicates. Relative amounts of mRNA were determined by comparing the samples to a standard curve, and expressed relatively to ribosomal RNA Rns18s.

For comparing longitudinal PER2::LUC recordings to the actual PER2::LUC protein levels (longitudinal versus acute luciferase assays), synchronised WT and CKO cells (cultured in absence of luciferin) were harvested every hour (in triplicate) from 16 hours up to 64 hours after media change, while co-cultures were recorded for bioluminescence in presence of luciferin. Luciferase activity in acute assays was determined as detailed in SI.

For assaying the interaction between BMAL1 and PER2::LUC, synchronised cells were harvested directly from temperature cycles at the expected peak of PER2::LUC expression (4 hours after change to 32°C) and BMAL1 was precipitated as described in SI. PER2::LUC co-immunoprecipitation was measured in a luciferase assay by mixing the BMAL1-loaded beads in luciferase assay buffer (15 mM MgSO4, 30 mM HEPES, 300 μM luciferin, 1 mM ATP, 10 mM 2-mercaptoethanol) and measuring luciferase activity in a Berthold platereader. The results were corrected for input and plotted relatively to the WT IgG pulldown.

To study the interaction of BMAL1 with S6K and eIF4, cells were synchronised by a 2-hour dexamethasone pulse, after which they were changed into normal growth medium. 12 and 24 hours after the medium change, BMAL1 immunoprecipitation was executed as described in SI. Samples were analysed by Western blot for presence of BMAL1, S6K and eIF4 (cell signalling, resp. #2708 and #2013)

### *Drosophila* experiments

All fly strains were kept in standard cornmeal food under 12 h:12 h LD cycles at constant 25°C (LD cycles). The following control strains were included in the experiments: *per01, Canton S,* and *w1118*. The generation of *Tim_Out_*flies, crossings with XLG-luc flies (Veleri et al., 2003), and details of recordings are described in SI. In short, three to seven days-old flies were entrained for three day LD cycles before being loaded individually into the wells of a microtiter plate containing the food-luciferin substrate (15mM luciferin). Recordings were performed under constant darkness at 26oC over seven days. Bioluminescence from each fly was background subtracted, summed into 2-hour bins, then detrended using a 24-hour moving average. Rhythmicity of averaged traces was tested using the RAIN algorithm (Thaben and Westermark, 2014) using 4-hour binned traces from 48 till 96 hours. Normalised and detrended Single fly traces were manually divided over three categories: “Robustly rhythmic”, “Poorly rhythmic”, and “Arrhythmic” according to examples in Figure S2B: traces with clear ~24hr rhythms over 4 cycles were termed “robustly rhythmic”, traces with lower amplitude but overt rhythms over 3 cycles “poorly rhythmic” and traces with <3 overt peaks were termed “arrhythmic”.

### Analysis

All analyses were performed in Graphpad Prism versions 7 and 8. Where indicated, data was detrended using moving average subtraction, where temporal window of the moving average was refined iteratively until it matched with the period of oscillation derived as follows. Period analysis was performed either manually, or by least-square fitting to a circadian damped sine wave with a linear baseline:

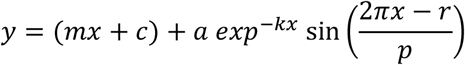

Where *m* is the gradient of the baseline, *c* is the *y* offset, *k* describes the rate of dampening, *a* the amplitude, *r* the phase and *p* the period. Reported p-values for the curve fit are those produced by the comparison of fits functions in Prism 8, where the null hypothesis was a straight line (y = mx + c), i.e., change over time but with no oscillatory component. The simpler model was preferred unless the sine wave fit produced a better fit with p<0.05.

For the mathematical model in 4E we used assumed that PER2::LUC translation at time (t) is a function of *Per2::Luc* mRNA abundance, corrected for the changes we observed for global translation rate over time; and that PER2::LUC degradation rate follows one-phase exponential decay kinetics where the decay constant is defined by a sine wave with 24-hour periodicity, with the amplitude, phase and other parameters being derived entirely from experimental measurements. See SI for details.

## Supporting information

Suppl data, discussion and methods

## Acknowledgements

We thank biomedical technical staff at Medical Research Council (MRC) Ares facility and LMB facilities for assistance, G.T. van der Horst and J.S. Takahashi for sharing rodent models, M.H. Hastings and E.S. Maywood for providing reagents and input, K. Lamia for providing reagents, and P. Crosby, D.S. Tourigny, J.E.C. Jepson, C.P. Kyriacou, H.R Pelham for valuable discussion. MP was supported by the Dutch Cancer Foundation (KWF, BUIT-2014-6637) and EMBO (ALTF-654-2014). JON was supported by the Medical Research Council (MC_UP_1201/4) and the Wellcome Trust (093734/Z/10/Z). NP and RF were supported by the Deutsche Forschungsgemeinschaft FKZ (Pe1798/2-1). AS and CPS were supported by the National Institutes of Health (GM118102).

## Author contributions

MP and JON designed the study, analysed the data and wrote the manuscript; AZ and NR performed mouse behavioural studies; MP, DW, ES, NH, KF and JON performed cell experiments; CS and AS generated MEF cell lines; ME performed SCN experiments; KC, RF, NP and JON performed fly experiments; JC performed tissue collection and husbandry; All authors commented on the manuscript.

## List of abbreviations

CCGs: Clock controlled genes
CHX: Cycloheximide
CK1: Casein kinase 1
CKO: CRY Knock out
CPKO: CRY PER Knock out
CPS: Counts per second
CP 20s: Counts per 20 seconds
CRY: Cryptochrome
GSK3: Glycogen synthase kinase 3
LL>DD: 12h:12h Light:light to dark:dark transition
MAF: Mouse adult fibroblast
MEF: Mouse embryonic fibroblast
PER: Period
RLU: Relative light units
SCN: Suprachiasmatic nucleus
SD: Standard deviation
SEM: Standard error of the mean
*Tim_out_*: Timeless knockout
TTFL: Transcriptional translational feedback loop
WCL: Whole cell lysates
WT: Wild type

