## Supplementary material for "CRYPTOCHROMES confer robustness, not rhythmicity, to circadian timekeeping": Suppl data, discussion and methods

FIGURE S1

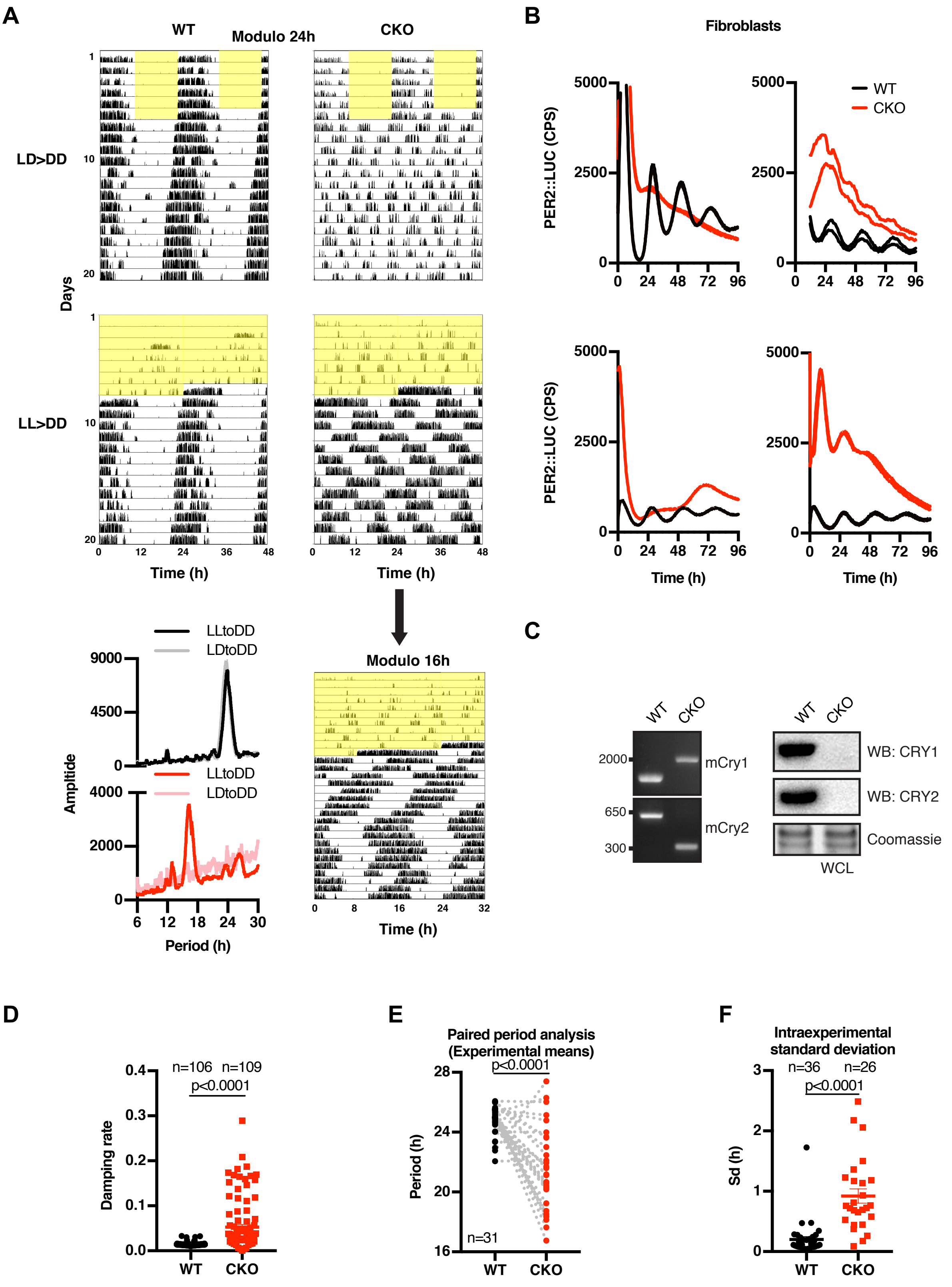
A
WT
Modulo 24h
CKO
LD>DD
LL>DD
Days
Time (h)
Amplitide
LLtoDD
LDtoDD
Period (h)
Modulo 16h
B
Fibroblasts
PER2::LUC (CPS)
WT
CKO
Time (h)
C
mCry1
mCry2
WB: CRY1
WB: CRY2
Coomassie
WCL
D
Damping rate
n=106 n=109
p<0.0001
E
Paired period analysis
(Experimental means)
Period (h)
n=31
F
Intraexperimental
standard deviation
Sd (h)
n=36 n=26

**FIGURE S1 CRY-independent circadian timekeeping occurs cell-autonomously**

(A) Representative double-plotted actograms showing wheel-running activity of WT and CKO mice during 12h:12h light:dark (LD) cycles (yellow shading indicating lights on) (top) or during constant light (LL) (bottom) and thereafter in constant darkness (DD). Top four figures have same x-axis (modulo 24 hours). Rhythmic behaviour of CKO mice in LL>DD condition becomes clear when plotting the data in 16-hour modulo (i.e. x-axis being 32 hours).

(B) Examples of independent bioluminescence recordings of PER2::LUC expression in CRY-deficient fibroblasts showing variability in shape and baseline of rhythmic CKO traces. Two representative traces are shown per experiment. Stringent entrainment, e.g. with temperature cycles or dexamethasone, increases the likelihood of observing rhythmicity, but only in approximately 30% of the experiments did we observe clearly rhythmic expression of PER2::LUC over 3 cycles. Despite our best efforts, over many years, we were unable to identify a set of entrainment and recording conditions that consistently produced CKO PER2::LUC rhythms and were forced to conclude more variables were in play than we were adequately able to control for.

(C) Genotyping CKO fibroblasts used throughout this study. Left: PCR genotyping shows the expected pattern of CRY1 and CRY2 knockout. Right: Western blot analysis of whole cell lysates (WCL) and probed with antibodies against CRY1 and CRY2.

(D) Interexperimental comparison of PER2::LUC periods in WT vs CKO fibroblasts. Paired comparison of period means of experiments used for Figure 1F where CKO traces were rhythmic. P-value were calculated in a paired t test.

(E) Intraexperimental standard deviations (i.e. between replicates) were calculated for all experiments with >3 rhythmic traces. P-value was calculated in an unpaired t test.

(F) Damping rates of all individual detrended traces of example experiments shown in Figure S1B were calculated by damped sine wave fitting. P-value was calculated in an unpaired t test with Welch correction.

#### FIGURE S2

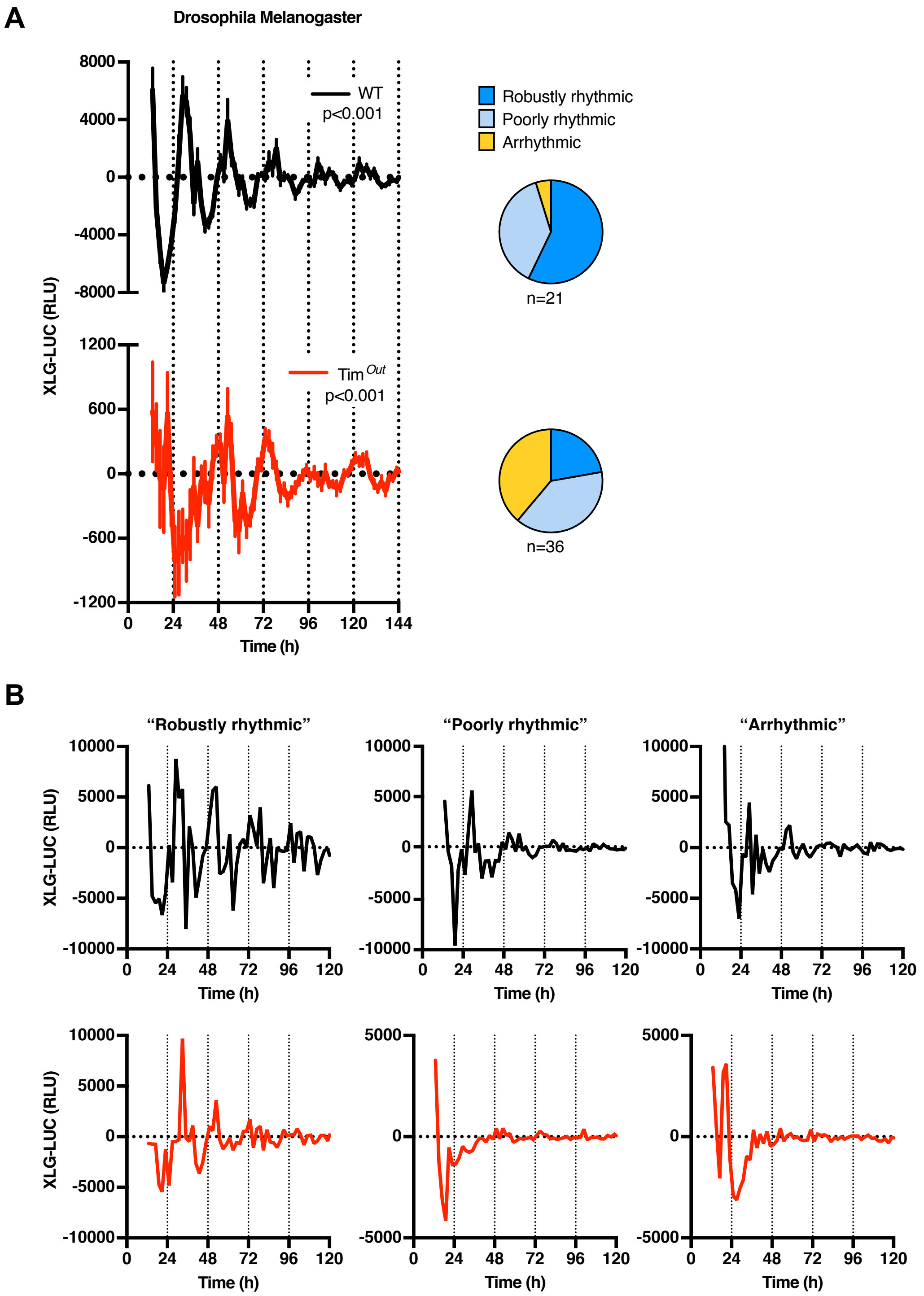

A
Drosophila Melanogaster
8000
4000
0
-4000
-8000
1200
600
0
-600
-1200
XLG-LUC (RLU)
WT
p<0.001
Tim $^{Out}$
p<0.001
0
24
48
72
96
120
144
Time (h)
Robustly rhythmic
Poorly rhythmic
Arrhythmic
n=21
n=36
B
"Robustly rhythmic"
"Poorly rhythmic"
"Arrhythmic"
10000
5000
0
-5000
-10000
XLG-LUC (RLU)
0
24
48
72
96
120
Time (h)
5000
0
-5000

**FIGURE S2. Timeless-independent protein rhythms in Drosophila Melanogaster**

(A) Normalised and detrended bioluminescence recording of the XLG-luciferase reporter (XLG-LUC; equivalent of PER2::LUC) expressed in WT and Timeless knockout ($Tim_{out}$) flies under constant darkness (detrended means ±SEM; WT n=21, $Tim_{out}$ n=36). Note the difference in Y-axis scaling. P-values report circadian rhythmicity (RAIN). Traces were manually divided over three categories: “Robustly rhythmic”, “Poorly rhythmic” and “Arrhythmic” and their distribution shown in the accompanying pie chart (right).

(B) Example of single fly recordings manually divided over three categories.

FIGURE S3

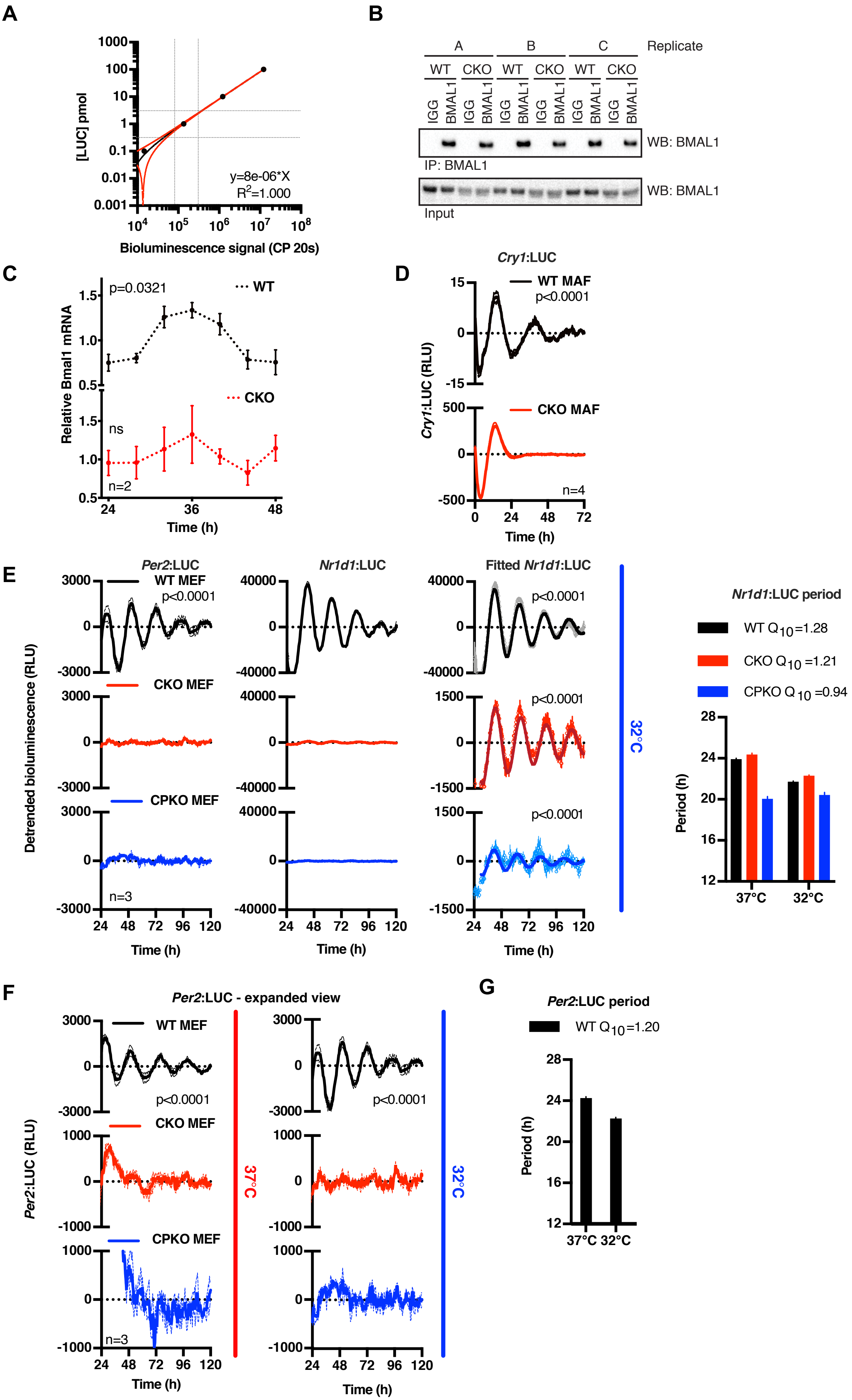

A
[LUC] pmol
y=8e-06*X
R2=1.000
Bioluminescence signal (CP 20s)
B
Replicate
A
B
C
WT
CKO
IGG
BMAL1
WB: BMAL1
IP: BMAL1
Input
C
p=0.0321
WT
CKO
Relative Bmal1 mRNA
ns
n=2
Time (h)
D
Cry1:LUC
WT MAF
p<0.0001
CKO MAF
Cry1:LUC (RLU)
n=4
Time (h)
E
Per2:LUC
Nr1d1:LUC
Fitted Nr1d1:LUC
WT MEF
CKO MEF
CPKO MEF
p<0.0001
Detrended bioluminescence (RLU)
n=3
Time (h)
32°C
Nr1d1:LUC period
WT Q10=1.28
CKO Q10 =1.21
CPKO Q10 =0.94
Period (h)
37°C
32°C
F
Per2:LUC - expanded view
WT MEF
CKO MEF
CPKO MEF
p<0.0001
Per2:LUC (RLU)
n=3
37°C
32°C
Time (h)
G
Per2:LUC period
WT Q10=1.20
Period (h)
37°C
32°C

**FIGURE S3 CRY-independent rhythms are regulated post-transcriptionally**

(A) Standard curve of recombinant luciferase that was used to determine the number of PER2::LUC molecules. Known concentrations of recombinant luciferase were spiked into (non-luciferase containing) cell lysate to reproduce experimental conditions, and the luciferase signal was measured with a 20 second integration time (CP 20s; counts per 20 seconds). Data were fitted with a straight line (red line, ± 95% CI). The grey dotted lines indicate the (linear) area of the curve used to determine the number of PER2::LUC molecules of the experiment shown in Figure 3A.

(B) Western blot analysis of BMAL1 co-immunoprecipitation samples shown in Figure 3B. BMAL1 or control (IgG) pulldowns were performed at the peak of PER2::LUC expression (determined in parallel PER2::LUC recordings) in 3 technical replicates (A-C).

(C) *Bmal1* mRNA levels were determined by qPCR over one circadian period (n=2, mean ±SEM). The WT timeseries were preferentially fit with a circadian damped sine wave compared with a straight line (p=0.0321), but not the CKO timeseries (ns). PER2::LUC co-recording from parallel cultures are depicted in Figure 3C.

(D) Detrended bioluminescence data of transcriptional reporter *Cry1*:LUC in WT and CKO mouse adult fibroblasts (MAFs) (n=4, mean ±SEM). WT traces fit circadian damped sine wave over straight line (p<0.0001), whereas no sine wave could be fit to CKO traces (no p-value).

(E) Detrended *Per2* and *Nr1d1* promoter activity in WT, CKO and quadruple *Cry1/2-Per1/2* knockout (CPKO) mouse embryonic fibroblasts (MEFs) recorded at 32°C. WT *Per2* and all *Nr1d1* traces were fit with a circadian damped sine wave over straight line (p<0.0001), whereas no sine wave could be fit to the other traces (no p-value). Period analysis shows that *Nr1d1* promoter oscillations are temperature compensated.

(F) Expanded view of *Per2*:LUC recordings to show no circadian oscillations of *Per2* promoter activity are detected in C(P)KO MEFs. Period analysis shows that also WT *Per2* promoter oscillations are temperature compensated, as expected.

### FIGURE S4

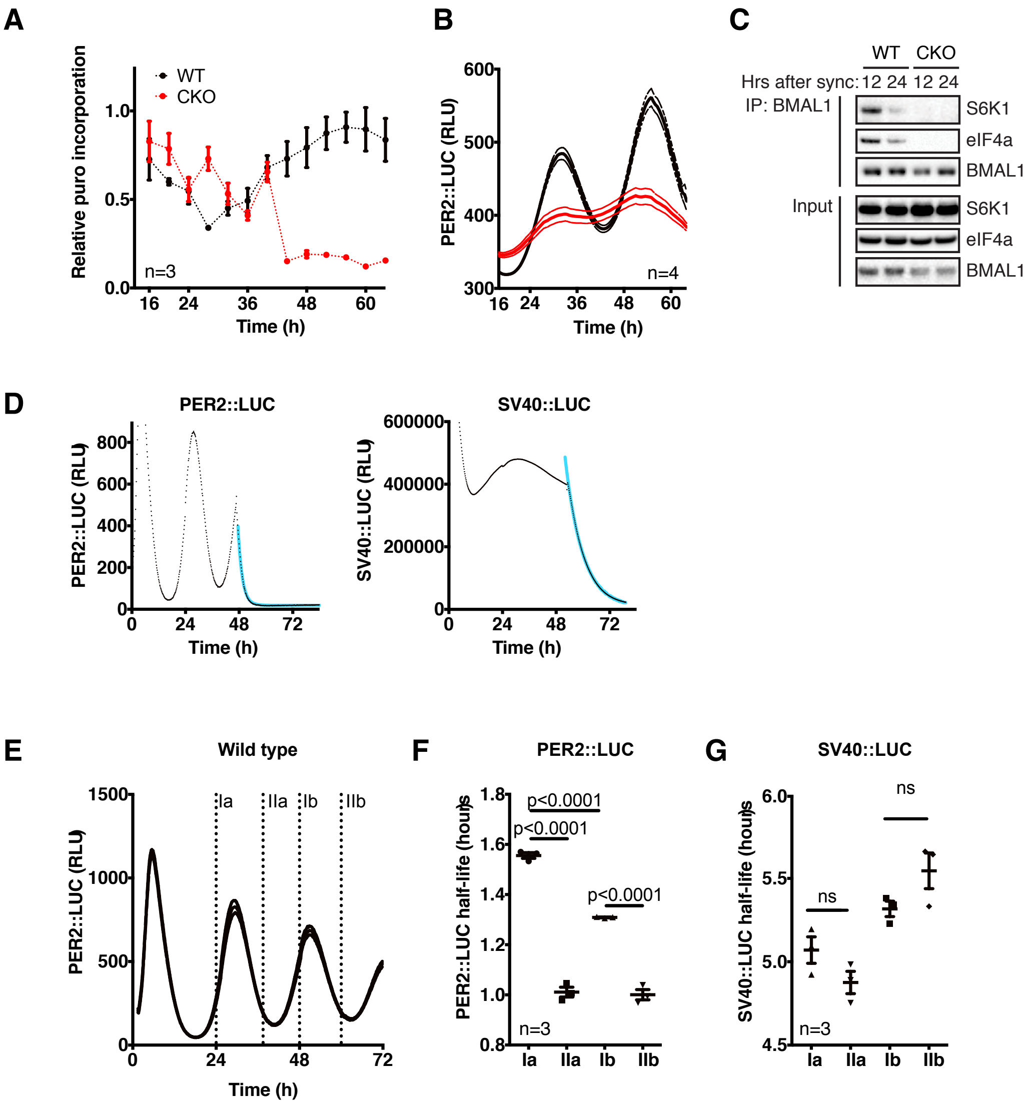
A
Relative puro incorporation
WT
CKO
n=3
Time (h)
B
PER2::LUC (RLU)
n=4
Time (h)
C
WT
CKO
Hrs after sync: 12 24 12 24
IP: BMAL1
S6K1
eIF4a
BMAL1
Input
S6K1
eIF4a
BMAL1
D
PER2::LUC
PER2::LUC (RLU)
Time (h)
SV40::LUC
SV40::LUC (RLU)
Time (h)
E
Wild type
PER2::LUC (RLU)
Ia
IIa
Ib
IIb
Time (h)
F
PER2::LUC
PER2::LUC half-life (hours)
p<0.0001
p<0.0001
p<0.0001
n=3
Ia
IIa
Ib
IIb
G
SV40::LUC
SV40::LUC half-life (hours)
ns
ns
n=3
Ia
IIa
Ib
IIb

**FIGURE S4 PER2::LUC stability oscillates in CRY-deficient cells**

(A) WT and CKO cells were assayed for puromycin incorporation over two circadian cycles. Cells were synchronised by temperature cycles and dexamethasone, and harvested every 3 hours after a 10 minute puromycin pulse (10 µg/mL). Incorporation was measured by Western blotting with an anti-puromycin antibody. Western blots were quantified and corrected for total protein loading (coomassie staining). Mean (n=3) ±SEM.

(B) Bioluminescence co-recording of puromycin labelling time course shows circadian PER2::LUC expression in both genotypes. Mean (n=3) ±SEM.

(C) Western blot analysis of BMAL1 immunoprecipitation with antibodies specific for S6K, eIF4a and BMAL1. Cells were harvested 12 or 24 hours after dexamethasone synchronisation and BMAL1 was immunoprecipitated.

(D) Example of a bioluminescence recording of WT PER2::LUC (left) or SV40::LUC (right) cells pulsed with 10 µM CHX after 46 hours of recording. The resulting raw data (symbols) were fitted with a one-phase decay curve (blue line).

(E) Timing of CHX pulses (labelled I-II a (cycle 1) and b (cycle 2)), plotted on bioluminescence trace of WT PER2::LUC control cells, corresponding to data presented in Figure S4F and G.

(F) PER2::LUC half-life at different phases in the circadian cycle in WT cells. Half-life was quantified by one-phase decay line-fitting of bioluminescence traces from CHX pulsed cells.

(G) SV40::LUC half-life at different phases in the circadian cycle in fibroblasts.

### FIGURE S5

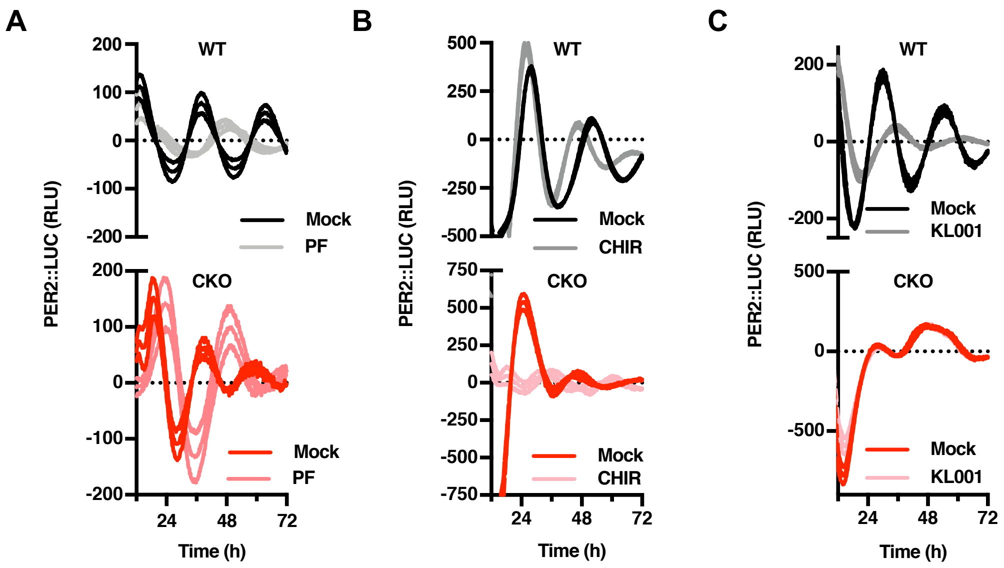

A
B
C
WT
CKO
PER2::LUC (RLU)
Time (h)
Mock
PF
CHIR
KL001
200
100
0
-100
-200
500
250
-250
-500
750
-750
24
48
72

**FIGURE S5. A role for CK1 and GSK3 in the cytoplasmic oscillator**

(A) Bioluminescence recordings of WT and CKO PER2::LUC cells in presence or absence of CK1δ/ε inhibitor PF670462 (0.3 µM), as quantified in Figure 5A (n=3, detrended mean ±SEM).

(B) As in (A), GSK3 inhibitor CHIR99021 (5 µM; CHIR).

(C) As in (A), in presence of CRY inhibitor KL001 (1 µM).

**Supplemental information, Putker et al.**

#### Supplementary discussion

Regarding previously reported CKO PER2::LUC arrhythmicity

A previous report suggests that no circadian bioluminescence rhythm in PER2::LUC activity is expressed by any tissues or cells derived from CKO mice (Liu et al., 2007). Besides our own finding, this earlier conclusion has already been contradicted for neonatal SCN slices by two independent labs (Maywood et al., 2011; Ono et al., 2013), and so it seems likely that some methodological difference must account for differences between their observations and those from other labs using the same genetic model. Critically, the CKO rhythms we observe conform to the formal definition of circadian rhythms: oscillations with (about) daily frequency, that are temperature-compensated and whose phase is sensitive to external stimuli (Pittendrigh, 1960).

Regarding the proposed model of coupled oscillators

We think the simplest interpretation of our findings entails an underlying, evolutionarily-conserved post-translational timekeeping mechanism: a "cytoscillator". This cytoscillator confers 24-hour periodicity upon the activity and stability of PER2, and most likely to other clock protein transcription factors as well (Figure S5C). We suggest that in wild type cells, low amplitude, cytoscillator-driven circadian cycles of clock protein activity are coupled with, reinforced and amplified by a damped TTFL-based relaxation oscillation of stochastic frequency, resulting in high-amplitude, sustained circadian rhythms in both clock and clock-controlled gene expression.

The model is attractive for several reasons. First, it may explain the discrepancy between SCN and behavioural studies in CKO mice, in that residual timekeeping can be observed in cultured SCN PER2::LUC activity, whereas behavioural rhythmicity is not observed in constant darkness following standard 12h:12h light:dark entrainment, but is expressed under specific non-standard conditions (Iijima et al., 2005; Ono et al., 2013). Considering the CKO cellular clock's shorter intrinsic period, as well as the profound robustness conferred upon SCN timekeeping by interneuronal coupling (Welsh et al., 2010; Yamaguchi et al., 2003), it seems plausible that 24h cycles may simply lie outside the range of circadian entrainment for CKO SCN *in vivo*, similar to the *tau* mutant hamster and humans with familial advanced sleep phase syndrome (Meng et al., 2008; Ptáček et al., 2007). Whereas *ex vivo*, or following the strong synchronising cue imposed by transition from constant light to constant darkness (Chen et al., 2008), non-transcriptional cellular mechanisms are sufficient to impart circadian regulation to CKO neuronal activity that is amplified by neuropeptidergic $Ca^{2+}$/cAMP-signalling (O'Neill and Reddy, 2012), facilitating the same temporal consolidation of locomotor activity observed in wild type mice but with shorter period and less precision.

Second, whilst the evidence is indisputable that transcriptional feedback repression is critical for circadian co-ordination of global gene expression, physiology and behaviour, the evidence that these regulatory gene expression circuits are inherently possessed of approximately 24-hour rhythmicity is weak (reviewed in (Lakin-Thomas, 2006; Putker and O'Neill, 2016; Wong and O'Neill, 2018). Post-translational regulation of clock protein stability, activity and localisation, however, is already well-established as the primary determinant of the delay constants that allow the oscillation to persist with a period of about one day in all studied eukaryotic cells (Gallego and Virshup, 2007; van Ooijen et al., 2011; Top et al., 2018; Wong and O'Neill, 2018). We simply suggest that transcriptional feedback repression is not essential for circadian timekeeping *per se*, but amplifies the rhythms to increase robustness via hysteresis, when engaged, and also to confer tissue and cell-type specific functionality (Wong and O'Neill, 2018). Our paradigm here being the cell division cycle, where the essential timing mechanism is also post-translational, and persists in enucleated cells (Hara et al., 1980; Pomerening et al., 2005).

Third, there is no evidence that TTFL-mediated oscillations would not damp to a steady state without post-translational input (Wong and O'Neill, 2018). In contrast, there are several examples in the eukaryotic lineage, where circadian timekeeping persists in the absence of cycling gene expression (Lakin-Thomas, 2006; O'Neill and Reddy, 2011; O'Neill et al., 2011; Sweeney and Haxo, 1961). For example, the period of circadian rhythms in human cells and *Ostreococcus tauri* is regulated by CK1, both in the presence and absence of nascent transcription (Beale et al., 2019; O'Neill et al., 2011) similar to the rhythm reported by PER2::LUC in CKO cells we report here.

#### Supplementary experimental procedures

##### Entrainment protocols for cell experiments

Fibroblast bioluminescence recordings were performed with confluent, quiescent monolayers. Due to iterative refinement of the synchronisation protocol used to optimise rhythms in the CKO fibroblasts, several different entrainment protocols were used throughout the paper, with appropriate WT controls. We found the most effective method employed temperature entrainment cycles (12h 32°C – 12h 37°C) for at least five days, prior to incubation of cells with 100 nM dexamethasone for 2 h beginning at 4 h after the start of the warm phase, followed immediately by changing cells into the experimental recording medium. Despite our best efforts, over many years, we were unable to identify a set of entrainment and recording conditions that consistently produced CKO PER2::LUC rhythms and were forced to conclude more variables were in play than we were adequately able to control for. Those that we tested are as follows: seeding density, passage number, glucose concentration, serum concentration, amino acid concentration, time in culture, time in temperature cycles, period length of temperature cycles, pH buffer, growth factor (B27) concentration, co-culture with WT cells and media conditioning. A more detailed description can be found online: https://www.repository.cam.ac.uk/handle/1810/300610. To reduce variation in cell attachment as a source of experimental error we used a fibronectin coating in some experiments; this did not significantly affect subsequent bioluminescence rhythms.

##### PER2 molecule count

Cells were seeded in 10% DMEM and entrained by temperature cycles for three days after which they were shifted to constant 37°C. At the following estimated peak of PER2::LUC activity (as reported in a co-recording), cells were trypsinised and washed in PBS, after which 5 x10$_6$ cells were lysed in 100 mM potassium phosphate buffer pH 7.8, 1 mM EDTA, 100 mM 2-mercaptoethanol, 1% triton, 10% glycerol and protease inhibitors. Lysates were cleared by centrifugation and diluted tenfold in 15 mM $MgSO_4$, 30 mM HEPES, 300 µM luciferin and 1 mM ATP. Dilutions were staggered to correct for time lags between bioluminescence measurements. PER2::LUC activity was measured in a plate reader and the number of molecules was calculated by comparing the bioluminescence signal to a standard curve of QuantiLum® Recombinant Luciferase (Promega) which was spiked into (luciferase-free) cell lysate to control for factors in the lysate that might affect bioluminescence. Three technical replicates were measured in every experiment and the experiment was carried out three times. A representative experiment is shown.

**qPCR time course**

Cells were seeded in 10% DMEM and entrained by temperature cycles over five days. 24 hours before the first time point, media was changed to 1% air medium (1% serum, +1 mM luciferin), after which the cells were placed in constant conditions (37°C) and the recording begun. Cells were harvested in triplicate every four hours from 24 hours up to 48 hours after media change. Cells were harvested in RLT buffer according to protocol from the Qiagen RNeasy kit and samples were immediately snap frozen. mRNA isolation was performed according to the manufacturer's protocol (including DNase treatment), as was cDNA synthesis (Biorad iScript™ cDNA Synthesis Kit). qPCR was performed using SYBR® FAST qPCR Kit (KAPA biosystems) on a Prime Pro 48 Real-time qPCR machine (Bibby Scientific) (for primer and protocol details see below). Analysis involved three technical and three biological replicates. Relative amounts of mRNA were determined by comparing the samples to a standard curve, and expressed relative to ribosomal RNA Rns18s.

**Primers**

Primers used for genotyping CRY1/2:

| Gene name | Genotype | Primer | Temp |
|---|---|---|---|
| *Cry1* | WT | CAGGAGGAGAAACTGAGGCACT | 63°C |
| | CKO | TGAATGAACTGCAGGACGAG | |
| | WT/CKO | GTGTCTGGCTAAATGGTGG | |
| *Cry2* | WT | CCAGAGACGGGAAATGTTCTT | 57°C |
| | CKO | GAGATCAGCAGCCTCTGTTCC | |
| | WT/CKO | GCTTCATCCACATCGGTAACTC | |

Primers used for QPCR:

| Gene name | Forward primer | Reverse primer | Temp |
|---|---|---|---|
| *Rns18s* | CGCCGCTAGAGGTGAAATTC | TTGGCAAATGCTTTCGCTC | 58°C |
| *Bmal1* | ACGACATAGGACACCTCGCAGA | CGGGTTCATGAAACTGAACCATC | 55°C |
| *Per2* | CCTACAGCATGGAGCAGGTTGA | TTCCCAGAAACCAGGGACACA | 58°C |

**Puromycin labelling time course**

Wild type and CRY-deficient PER2::LUC cells were seeded in fibronectin-coated 6-well plates in 10% DMEM and temperature entrained for five days. 18 hours before harvest they were pulsed with dexamethasone (100 nM) for 2 hours, and changed into air medium (HEPES, with 1 mM luciferin), after which the bioluminescence recording commenced. Cells were harvested every 4 hours (three biological replicates per time point) directly from the recording device from 16 hours up to 64 hours after media change. Ten minutes before harvest, cells were placed on a 37°C heat-pad and pulsed with 10 µg/mL puromycin. Cells were washed in ice-cold PBS, 5 mM EDTA and 20 mM NEM, harvested in 50 mM Tris-HCl pH 7.4, 150 mM NaCl, 0.1% LDS, 1% Triton, 0.5 % NaDOC, 20mM NEM and protease inhibitors, with samples then being immediately frozen in liquid nitrogen. After thawing, samples were cleared by centrifugation and taken up in reducing LDS sample buffer for Western blot analysis. Puromycin incorporation was assayed by Western blotting with specific anti-puromycin antibody (PMY-2A4-2, Developmental studies hybridoma bank). The signal was corrected for total protein loading on coomassie blue staining.

**Acute PER2::LUC measurements**

WT and CKO cells PER2::LUC cells were seeded in absence of luciferin in fibronectin-coated 35 mm dishes and synchronised by temperature entrainment, dexamethasone pulse and change into air medium. Cells for parallel co-recordings were pre-incubated with 0.1 mM luciferin. Cells were harvested every hour (in triplicate) from the recording incubator from 16 hours up to 64 hours after media change. Cells were washed in ice-cold PBS, lysed in 100 mM potassium phosphate buffer pH 7.8, 1 mM EDTA, 7 mM 2-mercaptoethanol, 1% triton, 10% glycerol, 1 mM NaF, 1 mM $Na_3VO_4$ and protease inhibitors, then samples were immediately frozen in liquid nitrogen. After thawing, samples were cleared by centrifugation and used for acute luciferase assays. Bioluminescence activity in 10 µL sample was measured in triplicate in a Spark 10M microplate reader (Tecan) and initiated by injection of 90 µL1.5 mM $MgSO_4$, 30 mM HEPES, 300 µM luciferin and 1 mM ATP. Immediate reading obviated the need for differential times of incubation prior to measurement.

**Co-immunoprecipitation experiments**

For assaying the interaction between BMAL1 and PER2::LUC, cells were entrained in temperature cycles for 4 days and harvested directly from temperature cycles at the expected peak of PER2::LUC expression (4 hours after change to 32°C). Cells were washed in ice-cold PBS and lysed in 200 µL 50 mM Tris-HCl pH 7.5, 1% TX100, 10 mM $MgCl_2$, 100 mM NaCl, DNase (100 U/mL) and protease inhibitors. After incubation at 4°C for 10 minutes, lysates were passed through a 19 gauge needle to ensure complete lysis, cleared by centrifugation, and diluted in 1 mL ice-cold wash buffer (50 mM Tris-HCl pH 7.5, 0.1% TX100, 5mM EDTA,1.5 mM $MgCl_2$, 100 mM NaCl and protease inhibitors). Total lysate samples were taken and stored at 4°C for the duration of the experiment. BMAL1 was precipitated with goat-anti-BMAL1 (Santa Cruz, SC-8550) antibodies or control IgG (SC-2028) coupled to protein G agarose beads (Pierce 20398) while rocking for two hours at 4°C. Samples were washed 3 times with ice-cold wash buffer and once in minimal luciferase assay buffer (15 mM MgSO4, 30mM HEPES). PER2::LUC co-immunoprecipitation (co-IP) was measured in a luciferase assay by mixing the beads in 200 uL luciferase assay buffer (15 mM MgSO4, 30 mM HEPES, 300 µM luciferin, 1 mM ATP, 10 mM 2-mercaptoethanol) and measuring luciferase activity in a Berthold platereader. The results were corrected for input (5 µL of the total lysate sample) and plotted relatively to the WT IgG pulldown. After measurements, the beads were washed in wash buffer and taken up in sample buffer for Western blot analysis of BMAL1 pull-down efficiency (homemade rabbit-anti-BMAL1 antibody (Sládek et al., 2007)). To study the interaction of BMAL1 with S6K and eIF4, cells were entrained by a 2-hour dexamethasone pulse, after which they were changed into normal growth medium. 12 and 24 hours after the medium change, BMAL1 immunoprecipitation was executed as described above. After washing with wash buffer, samples were taken up in reducing LDS sample buffer and analysed by Western blot for presence of BMAL1, S6K and eIF4 (cell signalling, resp. #2708 and #2013)

##### ***Drosophila* experiments**

##### **Fly stocks and husbandry**

The XLG-luc construct contains a fusion between Period CDS and Luciferase under endogenous control of the *period* promoter and flanking regulatory regions, as described previously. The luciferase signal from the derivative fly strain, *y w;; XLG-luc:2/TM3,* faithfully reports the endogenous PERIOD protein rhythm (Veleri et al., 2003).

##### Generation of *tim*$_{Out}$ fly

*Timeless* knock-out fly lines (*tim*$_{KO}$) were generated by homologous recombination (Huang et al., 2009) and described in (Lamaze et al., 2017). The residual mini-*white* marker in the *tim*$_{KO}$ flies was floxed-out. The obtained knockout lines were denoted as *w*$_{1118}$; *tim*$_{Out}$. One derivative strain, *y w; tim*$_{Out}$*/CyO; XLG-luc:2/TM3*, was generated via standard balancer crossing.

##### Longitudinal XLG-luciferase recordings

To monitor circadian rhythm of PER::LUC activity, we crossed male *y w;; XLG-luc:2/TM3 and y w; tim*$_{Out}$*/CyO; XLG-luc:2/TM3* flies with two female control, *w*$_{1118}$ and *Canton S*, and to two clock mutants, *per*$_{01}$ and *w*$_{1118}$*; tim*$_{Out}$, respectively. The resultant F1 males: *per*$_{01}$*/Y*;+/+;*XLG-luc:2/*+, *w*$_{1118}$*/Y*;+/+;*XLG-luc:2/*+, +$_{Canton\ S}$*Y ;*+/+*;XLG-luc:2/*+, and *w*$_{1118}$*/Y;tim*$_{Out}$*/tim*$_{Out}$*;XLG-luc:2/*+ were studied.

Three to seven days old flies were then entrained for three day LD cycles before being loaded individually into the wells of a microtiter plate containing the food-luciferin substrate (15mM luciferin) where their movement was restricted by covering them with pierced plastic domes (Stanewsky et al., 1997). Recordings were performed under constant darkness at 26°C over seven days. Bioluminescence images were recorded with contiguous 5 min integrations over 7 days with camera settings. The plate was placed in the ALLIGATOR and recorded in parallel with the tube-based assay condition. Bioluminescence from each fly was background subtracted, summed into 2-hour bins, and then detrended using a 24-hour moving average. Since the per$_{01}$ and control XLG-luc flies produced very similar bioluminescence traces, for clarity only the *tim*$_{Out}$ and WT control data are shown.

**Mathematical modeling**

The mathematical model we used assumed that PER2::LUC translation at time (t) is a function of *Per2*::*Luc* mRNA abundance, corrected for the changes we observed for global translation rate over time; and that PER2::LUC degradation rate follows one-phase exponential decay kinetics where the decay constant is defined by a sine wave with 24-hour periodicity, with the amplitude, phase and other parameters being derived entirely from experimental measurements, as follows:

$P_t = P_{t-1} + S_t - D_t$

$S_t = 1000*R_t*T$

$D_t = P * (1- e_{-kt})$

$k_t = \ln(2)/((A(\sin((2\pi*t/24)+\varphi)))+H)$

t  = time in hours

$P_t$ is PER2::LUC protein abundance at t (from measured number of molecules/cell)

$S_t$ is total P translated in 1 h prior to t

$D_t$ is total P degradation in 1 h prior to t

$R_t$ is *Per2*::*Luc* mRNA abundance at t (interpolated from qRT-PCR measurements)

$T_t$ is translation rate at t (interpolated from puromycin incorporation assays)

$k_t$ is the exponential decay constant at t (from PER2::LUC half-life measurements)

A is amplitude of the rhythm in PER2::LUC half-life (from PER2::LUC half-life measurements)

$\varphi$ = initial phase (from observed phase of PER2::LUC relative to *Per2*::*Luc* mRNA level)

H = is mean PER2::LUC half-life (from PER2::LUC half-life measurements)
